# Defining focal neuroendocrine differentiation as a transcriptionally distinct form of prostate cancer pathology characterized by the expression of androgen receptors

**DOI:** 10.1101/2024.03.17.585125

**Authors:** Rosalia Quezada Urban, Shivakumar Keerthikumar, Ashlee Clark, Hong Wang, Belinda Phipson, Andrew Bakshi, Andrew Ryan, Heather Thorne, Renea A Taylor, Mitchell G Lawrence, Gail Risbridger, Roxanne Toivanen, David L Goode

## Abstract

**Background:** Men with neuroendocrine prostate cancer (NEPC) have a poor prognosis. NEPC is commonly diagnosed by immunohistochemical markers (CHGA, SYP and NCAM1) and genomic features (mutations in RB1, PTEN, TP53). But by pathology, NEPC tumours are variable, leading to a classification of NE subtypes such as small cell and large cell neuroendocrine carcinomas, focal neuroendocrine differentiation (Focal NED), and Amphicrine. We postulated the diversity observed in NEPC pathologies might arise from differences in transcriptional profiles and the aim of this study is to utilize single-cell RNA sequencing to define the transcriptional differences between NEPC subtype pathologies.

**Methods:** Gene expression profiles were obtained for 18,632 individual tumour cells from 9 patient-derived xenograft (PDX) models representing five distinct neuroendocrine pathologies of prostate cancer. Integration and clustering of cell-level data demarked transcriptionally distinct sub-populations of cells. Differential gene expression, gene set enrichment and transcriptional factor regulon analysis identified expression signatures unique to specific neuroendocrine pathologies. Copy-number estimated from expression data revealed the clonal structure of PDXs with mixed adenocarcinoma and neuroendocrine pathologies.

**Results:** Significant differences were observed in the transcriptional profiles of NEPC pathology subtypes. Focal NED cells maintain AR signaling, similar to the amphicrine subtype but different from small and large cell carcinomas. Cellular sub-populations enriched for expression of KRAS, IL2-STAT5 and TNF-signaling genes were found in focal NED and amphicrine pathologies, but not in small or large cell carcinomas. In contrast, sub-populations enriched for the YAP, Myc and E2F pathways were detected in small cell, large cell and amphicrine tumours, but not in focal NED cells. Each pathology showed unique patterns of master regulator activity as well, further implicating focal NED as a transcriptionally distinct entity. Based on copy number alterations within PDXs of mixed pathology, focal NED cells showed little clonal divergence from neighboring adenocarcinoma cells, whereas cells with small cell neuroendocrine pathology were clonally distinct.

**Conclusions:** Neuroendocrine prostate cancer subtypes can be identified by pathology and our study shows that transcriptional features identified by single-cell RNA-sequencing also distinguish neuroendocrine subtypes pathologies from each other. In particular, our data redefine focal neuroendocrine differentiation as a pathology expressing androgen receptors (AR), exhibiting its distinctive composition of transcriptionally unique sub-populations. These findings advocate for differences in the treatment of NEPC tumors, particularly those displaying focal NED.

## Introduction

Neuroendocrine prostate cancer (NEPC) is an aggressive subtype of prostate cancer diagnosed on the basis of immunohistochemistry (IHC) of canonical neuroendocrine cell surface markers such as chromogranin A (CHGA), synaptophysin (SYP), and CD56 (NCAM1) (Kannan et al, 2022). NEPC can arise via lineage plasticity under prolonged androgen deprivation (Beltran et al., 2016, Aggarwal et al., 2018), but it can also appear *de novo* at diagnosis (Epstein et al., 2014; Fine, 2018). No effective long-term treatments exist for NEPC and overall patient survival rates are very poor (Aggarwal et al., 2014; Aggarwal et al., 2018). NEPC is often associated with suppression of androgen receptor (AR) activity. Small cell and large cell neuroendocrine carcinomas are two prostate cancer pathologies that typically lack detectable AR signalling and are most often associated with NEPC (Epstein et al., 2014; Fine, 2018).

Additional neuroendocrine pathologies have been observed in prostate cancer (Aggarwal et al., 2014; Bellur et al., 2019; Epstein et al., 2014), defined by histology and morphology (Beltran et al., 2016). In contrast to small and large cell carcinoma, the amphicrine pathology is defined by strong co- expression of both AR activated and neuroendocrine genes (Epstein et al., 2014; Fine, 2018). Prostate adenocarcinoma with focal neuroendocrine differentiation (NED) displays small, scattered pockets of cells expressing neuroendocrine markers. Focal NED does not fully adhere to accepted definitions of NEPC (Epstein 2014; Fine 2018), and its influence on clinical outcomes remains uncertain (Kardoust Parizi et al., 2019). Mixed tumours containing both adenocarcinoma and small cell pathologies occur as well (Epstein 2014; Fine 2018).

To date, small and large cell pathologies have been much better represented in genomic and transcriptomic studies than other pathologies with neuroendocrine features. The molecular foundations and therapeutic implications of diversity among neuroendocrine pathologies in prostate cancer thus remain elusive, contributing to suboptimal patient outcomes (Beltran et al., 2011). Mutations to *RB1*, *PTEN*, *TP53*, as well as upregulation of N-MYC, SOX2, BRN2, and ONECUT2 are recurrent in NEPC (Beltran et al., 2011; Davies et al., 2020; Labrecque et al., 2019) but none are exclusive to any neuroendocrine pathology.

Single-cell RNA-sequencing enables discovery and expression profiling of transcriptionally distinct cell populations within tumours, offering a way to directly characterize rare, dispersed pathologies such as focal NED. Single-cell RNA-sequencing studies of NEPC remain limited in scope but have uncovered substantial intra-tumoural heterogeneity at the transcriptional level. Key insights include evidence NEPC arises from luminal-like cells (Dong et al., 2020), elucidation of the roles of RB1, N- Myc and E2F in neuroendocrine trans-differentiation (Brady et al., 2021) and the resolution of hierarchies of transcription factors networks (Wang et al., 2022).

To explore how transcriptional intra-tumoural heterogeneity contributes to diversity of neuroendocrine pathologies in prostate cancer, we performed single-cell RNA-sequencing on nine patient-derived xenograft (PDX) models covering five distinct pathologies of NEPC. Variation in both the type and frequencies of transcriptionally distinct cellular sub-populations was seen between PDXs of different pathologies. Focal NED cells displayed unexpected co-expression of AR signalling and NE markers as well as differential patterns on oncogenic pathway expression, marking focal NED as a distinct molecular entity within the landscape of NEPC.

## Methods

### Patient Derived Xenografts

Patient derived xenografts (PDXs) were acquired from the Melbourne URological Research ALliance (MURAL). The PDXs lines are maintained in compliance with Monash University animal ethics approval (MARP 2014/085). The maintenance of the serially transplantable PDXs have been described previously (Risbridger et al., 2021). Briefly, PDXs are maintained by sub-renal or sub- cutaneous grafting into 6-8-week-old immunocompromised male NSG mice. The NSG mice are supplemented with 5mm testosterone implants for mixed or amphicrine pathologies, or surgically castrated mice for pure NE/AR null pathologies.

### Dissociation of Patient Derived Xenografts

PDXs were harvested from host mice and cut into 2 X 2 mm pieces using a scalpel. Tumour pieces were digested in 15mL RPMI, pencilling/streptomycin containing 13 U LiberaseTM (Sigma) and 3mg DNase (Roche), for 1 hour at 37C. Samples were disrupted with a pipette every 30 minutes during incubation to ensure suspension of cells. After cells were spun at 5 minutes at 1000rpm, red blood cells were lysed using Red cell Lysis buffer (Sigma) for 1 minutes. Red cell lysis was stopped with RPMI with 10% FBS. Cells were then resuspended in PBS, 1mM CaCl2, with 2% FBS and underwent negative selection for viable cells using the Easy Sep Dead Cell Removal kit (Miltenyi), according to the manufacturer’s protocol. After selection, cells passed through a 30uM cell strainer (Miltenyi). Cell viability was assessed using Trypan Blue. Samples with cell viability >80% were resuspended in PBS containing 2% BSA and proceeded to single cell analysis.

### Single cell RNA-Sequencing library preparation

scRNA-Seq was done on dissociated PDXs using the 10X Genomics Chromium Single Cell 3′ Library & Gel bead Kit V3.0, per the manufacturer’s instructions (CG000183 Rev C). Briefly, 5000 PDX cells were utilised per sample as input. By encapsulating cells in microfluidic droplets, around 4000 single-cell transcriptomes were recovered per sample. After reverse transcription, barcoded cDNA was purified with SILANE Dynabeads and amplified through 11 cycles of PCR. On an Agilent Bioanalyzer High Sensitivity chip, SPRIselect purification was performed to quantify the fragment size and concentration of the amplified cDNA. Libraries were sequenced on an Illumina NovaSeq6000 using paired-end reads of 151 base pairs.

### Expression quantification for individual cells

Paired FASTQ files were aligned to the indexed GRCh38 human and mm10 mouse reference genome using XenoCell v1.0 (Cheloni et al., 2021). Further, the human specific cells were extracted using a maximum 10% threshold of mouse specific reads in XenoCell. The filtered human specific paired reads were quantified using Alevin (Salmon Software v1.2.1) tool (Srivastava et al., 2019) by aligning against the GRCh38 transcriptome. The quantified matrix file was further imported into Seurat v3.2.0 (Hao et al., 2023) in R V4.2.0 (R Core Team, 2023) for all the downstream analysis.

### Identification and profiling of transcriptional sub-populations [within each PDX]

Quality control, implemented using Seurat (v 3.2.0), aimed to exclude outlier cells with low-quality features. Standardized filtering criteria were then applied to all samples, involving the exclusion of cells expressing fewer than 50 genes, those with fewer than 1000 genes, and sample-specific variations, including a high mitochondrial transcript fraction (range 25-30%) and a high transcript count (range 40,000 – 100,000) (see Supplementary Table 1).

### Cell Cycle Phase Identification

To ascertain the cell cycle phase of individual cells, the “CellCycleScoring” function was employed. Canonical cell cycle markers (Kowalczyk et al., 2015), were incorporated into Seurat, with a specific focus on features associated with the G2/M phase and markers indicative of the S phase. These elements were utilized as essential input parameters for the “CellCycleScoring” function, which effectively scored and classified each cell into distinct phases, namely “S,” “G2/M,” and “G1.”

### Normalization, Scaling, and Feature Identification

For the normalization, scaling, and identification of high variable features, the SCTransform function was utilized. This normalization method relies on Pearson residuals derived from “regularized negative binomial regression,” (Hafemeister et al., 2019) employing cellular sequencing depth as a covariate within a generalized linear model (GLM). Default parameters were applied. Subsequently, Principal Component Analysis (PCA) was executed using the top 3000 most highly variable features. The determination of the appropriate dimension was facilitated by an Elbow plot in subsequent steps.

### Clustering and Visualization

To initiate the clustering process, the “FindNeighbours” function in Seurat facilitated the construction of a Nearest-neighbour graph, utilizing default settings. Dimensions were then selected based on individual object (sample) characteristics. The “FindClusters” function employed the shared nearest neighbour (SNN) approach to identify distinct clusters of cells, with default parameters utilized, and the resolution determined per sample. Visualization of clustered cells was achieved through the Uniform Manifold Approximation and Projection (UMAP) dimensional reduction technique using the “RunUMAP” function, employing default settings and the previously selected dimensions.

### Optimal Cluster Determination

To ascertain the optimal number of clusters, the clustree function from the R package ClusterTree (Zappia et al., 2018) as employed. This function elucidates the division of clusters as resolution increases, providing valuable insights. The number of clusters was determined through the construction of a clustering tree spanning resolutions from zero to 1 in increments of 0.1. Optimal resolutions for each sample were carefully chosen. Subsequently, a set of resolutions was selected and subjected to testing. Resolution testing involved a comprehensive analysis of differentially expressed markers per cluster at each resolution. Resolutions with marker overlap in multiple clusters were systematically discarded to refine the determination of the optimal number of clusters. This meticulous approach ensured the robustness of the clustering outcomes.

### Differential Gene Expression Analysis

Identification of marker genes per cluster was conducted using the FindAllMarkers function within Seurat, employing a negative binomial test. Parameters included a log fold change threshold of 0.25 and a minimum fraction of 0.25 for genes detected in either of the two populations. Expression profiles of selected genes were visualized on a logarithmic scale, facilitating a comprehensive assessment. The difference in expression fraction between the two groups was calculated to discern distinctive patterns. The top five differentially expressed genes were chosen based on the highest difference and the highest average log fold change, thereby ensuring robust selection criteria. Manual curation was applied to select unique markers with pronounced expression patterns.

For gene set enrichment analysis, the log fold change threshold was adjusted to 0, and the minimum fraction of genes detected in either of the two populations was set to 0. This modification was crucial for enhancing sensitivity and specificity in identifying enriched gene sets associated with the differential expression patterns observed.

### Cancer Signature Analysis

To examine the expression of cancer signatures, the CancerSEA database (Yuan et al. 2019) was obtained. All gene sets from the database were downloaded and subsequently utilized to compute scores per cell using the “AddModuleScore” function within Seurat. Visualization of the proportion and expression patterns of the top five differentially expressed markers and signatures per cluster was accomplished using the “Dotplot” function. This approach provided a comprehensive and visual representation of the distinctive features and signatures associated with cancer expression patterns within individual clusters.

### Gene Set Enrichment Analysis

To elucidate enriched pathways across clusters, a comprehensive gene set enrichment analysis (GSEA) was executed. The “msigdbr” package, providing Molecular Signatures Database (MSigDB) (LIberzon et al., 2015) gene sets commonly utilized in GSEA, was employed alongside the “fgsea” R package for the analysis (Korotkevich et al., 2021). All genes differentially expressed in each cluster were pre-ranked based on the highest difference. The “fgseaMultilevel” function from the “fgsea” R package was deployed to conduct the enrichment analysis, with default settings employed, except for “nPermSimple,” which was set to 10000 to enhance the accuracy of P-value estimation. The utilized gene sets encompassed Hallmarks (H), Oncogenic (C6), and KEGG (CP), offering a comprehensive exploration of the pathways enriched within the distinct clusters.

### Integration: Cluster Similarity Spectrum (CSS) in Simspec

Integration of single-cell data using the Cluster Similarity Spectrum (CSS) algorithm in the Simspec package requires a Seurat object (He et al., 2020). Prior to integration, the data underwent preprocessing in Seurat, involving normalization, identification of variable features, data scaling, PCA, and dimensional reduction using UMAP. The “cluster_sim_spectrum” function was employed for data integration, utilizing the Pearson correlation method and “corr_kernel” as the spectrum type. Cluster resolution was set at 0.3, and the label tag was defined as the sample name. Following integration, UMAP and PCA were run for dimensional reduction, using “css” and “css_pca” as the reduction types, respectively, with ten dimensions selected for each step. Subsequently, the “FindNeighbors” and “FindClusters” functions were applied to calculate clusters after integration, with a resolution set at 0.3 and 10 dimensions utilized.

### Quality Control After Integration

To evaluate the success of integration and discern technical and biological sources of variation, multiple factors were considered. Cell cycle phase, transcript counts, and mitochondrial and ribosomal percentages were visualized for technical sources using the feature plot function from Seurat. Mitochondrial and ribosomal percentages were computed using the “PercentageFeatureSet” function from Seurat. Biological variation was assessed through a differential gene expression (DGE) analysis using the “FindAllMarkers” function from Seurat. This comprehensive quality control step ensured a thorough examination of the integrated data, distinguishing between technical and biological factors contributing to variation.

### Downstream analysis for integrated dataset

Following integration, downstream analyses including differential gene expression (DGE) and gene set enrichment analysis were executed. DGE analysis was performed same as described above, utilizing the “FindAllMarkers” function from the Seurat package. For gene set enrichment analysis (GSEA), MSigDB datasets were employed. Similar to the previous GSEA analysis on individual samples, the “fgseaMultilevel” function was utilized for the enrichment analysis, employing default parameters.

### Co-Expression Analysis

Co-expression analysis was conducted using the “Featureplot” visualization function within Seurat, with the “blend” argument set to TRUE. This setting enabled the simultaneous visualization of two markers’ expression on each cell in the UMAP. The co-expression scale, ranging from 0 to 10, was established, where 0 represents the lowest and 10 the highest expression. A maximum cut-off value of q25 (quantile) was set to capture the minimum expression of markers. The “blend” threshold was set to 0.1, initiating the blending of selected colours from the weakest signal. The percentage of cells co-expressing selected markers was determined by fetching normalized counts for each marker and calculating the co-expression percentage across cells.

### Scoring activity of transcription factor regulons with SCENIC

The integrated R object file containing raw counts matrix file was loaded into R and gene regulatory networks was inferred using PySCENIC package (v0.11.2) (Aibar et al., 2017). Regulons for which >20 target genes were identified was used further and its activity was depicted in heatmap s.

### Identification of clonal sub-populations

Clonal sub-populations were defined by chromosomal arm level copy-number differences using Gaussian mixture models to identify regions of the genome where contiguous genes show consistent increased/decreased expression within subsets of cells in a single-cell RNA-sequencing data set, in a reference-free manner (Kinker et al., 2020). Code for our analysis was adapted from the module5_cna_subclones.R script available at https://github.com/gabrielakinker/CCLE_heterogeneity.

Libraries for whole-genome sequencing (WGS) were prepared using the TruSeq DNA Nano High Throughput kit (Illumina) and sequenced as 150bp paired-end reads on a NovaSeq 6000 (Illumina). Reads from PDXs were aligned to hg19 (Ensembl Homo_sapiens.GRCh37.73.dna) and mm10 (Ensembl Mus_musculus.GRCm38.73.dna) using BWA MEM (v0.7.17), with duplicates marked by Picard (v2.17.3). Xenomapper (v1.0.1) (Wakefield, 2016) was used to identify reads mapping to hg19 only. The patient germline (blood) sample was aligned and processed in the same fashion but to the hg19 reference only.

PDX and germline BAM files were sorted with samtools (v1.9) and provided as input for clonality assessment based on copy-number alterations using the HATCHet algorithm (v0.1) (Zaccaria et al., 2020) with Gurobi Optimizer (v9.1.1, Linux 64-bit). Parameters were set as follows: mapQ=11, baseQ=11, snpQ=11, minCov=10, maxCov=300, binSize=100kb, with the sensitivity parameter (-l) set to 0.4.

## Results

### Establishing patient-derived models of neuroendocrine pathologies in prostate cancer

To better represent the heterogeneity of prostate cancer in the clinic, the Melbourne Urological Research Alliance (MURAL) established a collection of patient-derived xenografts (PDXs) spanning treatment-naïve primary prostate cancer to castration-resistant metastases (Risbridger et al., 2021).

This study focuses on nine MURAL PDXs with neuroendocrine features, including 8 published models (Risbridger et al., 2021) and a newly described PDX (470B). Each has undergone thorough histological assessment, along with genomic and transcriptomic profiling, to accurately annotate its pathology and confirm fidelity with the neuroendocrine phenotypes of the original donor patient (Risbridger et al. 2021).

The selected PDXs represent a variety of histopathologies, including adenocarcinoma with neuroendocrine differentiation (Focal NED; n=2), amphicrine carcinoma (n=1), mixed adenocarcinoma-small cell (n=1), small cell (n=2), large cell prostate cancer (n=2) and low-grade neuroendocrine carcinoma (n=1) (Figure 1). In each case, the histopathology of the PDX reflects the features of the original patient tumour.

**Figure 1:**
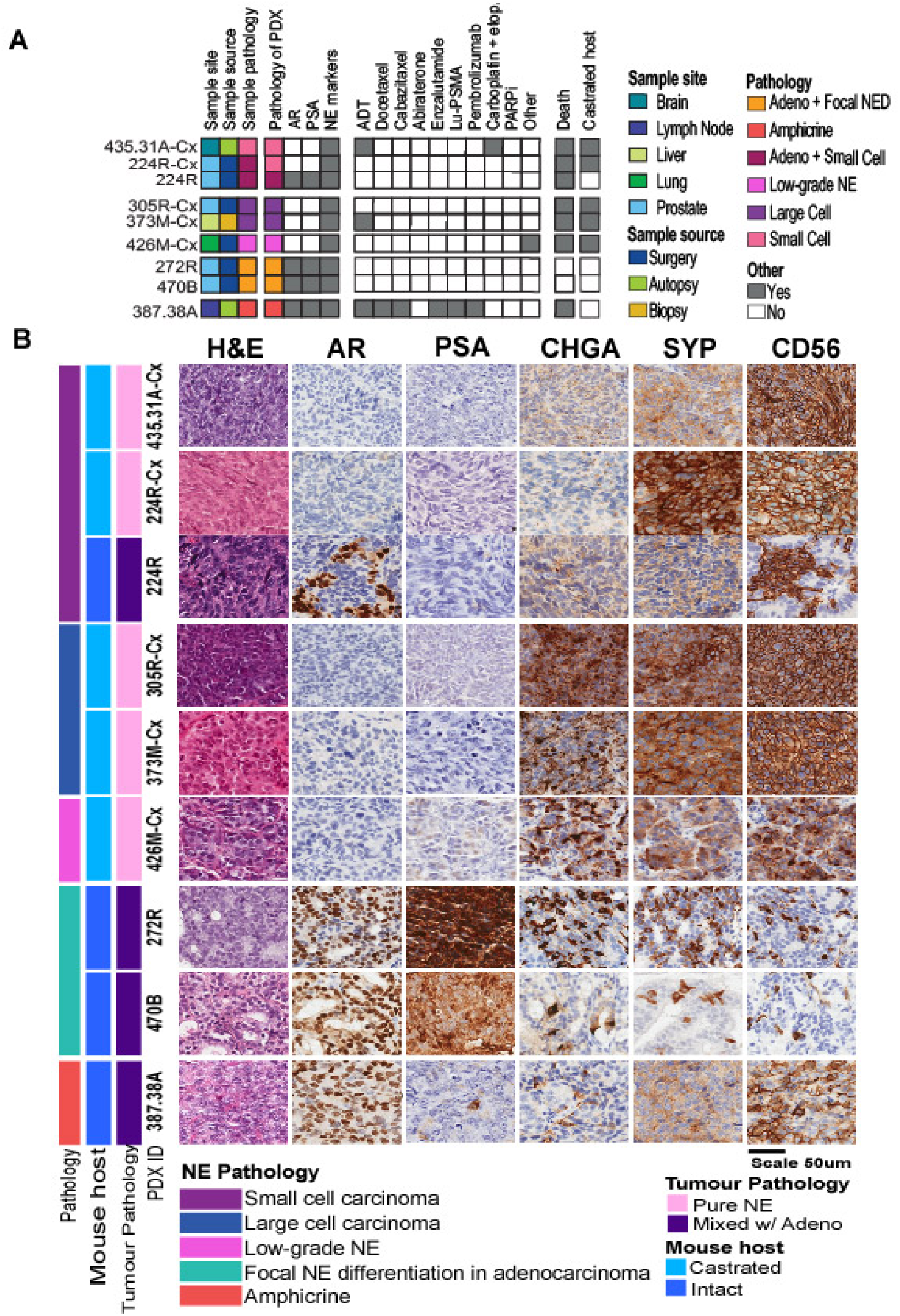
Diverse clinical and pathological landscape of MURAL PDXs with neuroendocrine features. (A) Clinical characteristics of the donor tumours used to establish of PDX models included in this study, the heatmap summarises the features of the patient samples, pathology of the PDXs, the patients’ treatment histories, collection method, follow-up, and whether the PDXs are maintained intact mice with testosterone implants or castrated mice. (B) Histopathology of PDX tumours, showing tissue morphology and staining for protein markers of adenocarcinoma (AR, PSA) and neuroendocrine (CHGA, SYP, CD56) Sidebar indicate assigned PDX tumour pathology and mouse host type.

Four PDXs originate from primary tumour samples donated at the time of radical prostatectomy from patients who had not received any systemic therapies (224R, 305R-Cx, 272R, 470B). The other four PDXs originate from metastases via biopsy, metastasectomy or from a rapid autopsy from patients with prior treatment, including ADT, androgen receptor signalling inhibitors, taxane chemotherapy, platinum chemotherapy, and Lu-PSMA (435.31A-Cx, 373M-Cx, 426M-Cx, 387.38A) (Figure 1A). Notably, patient 426M-Cx was diagnosed with de novo neuroendocrine prostate cancer at a very young age (before 30), while patient 470B had a germline *BRCA2* mutation.

All patient samples were initially grafted into immunocompromised mice with testosterone implants (Risbridger et al., 2021). Several PDXs continue to be grown under these conditions (224R, 272R, 387.38A, 470B). Other PDXs were subsequently regrafted in castrated host mice to simulate patients undergoing ADT (305R-Cx, 373M-Cx, 426M-Cx, 435.31A-Cx and 224R-Cx) [Table S1]. The tumour from patient 224 was maintained under both conditions, providing two PDX sublines. The PDX from testosterone-supplemented mice (224R) has mixed adenocarcinoma-small cell pathology, while the PDX from castrated mice has pure small cell (224R-Cx) pathology. Targeted exome sequencing revealed an abundance of alterations to *TP53*, *RB1* and *PTEN* in these PDXs, which is common in NEPC [Supp Fig S1]. Overall, these PDXs represent diverse forms of prostate cancer with neuroendocrine features.

### Prostate cancer cells with neuroendocrine pathology include a diverse array of transcriptional states

To analyse the heterogeneity of tumours with neuroendocrine features at single cell resolution, we obtained the transcriptional profiles of the nine PDXs using the 10X Genomics Chromium Single Cell 3′ sequencing chemistry (Methods). After removing mouse cells using Xenocell and iterative filtering of low-quality cells via Seurat (Methods) 1,202 – 7,796 cells were detected per PDX (mean: 2,659) [Supp Table S1]. The average number of genes detected per cell per PDX ranged from 2,829 – 8,037 (mean: 4,869.6). This demonstrates the robustness of our protocols for isolation and sequencing individual cells from prostate cancer PDXs.

Cells were clustered into subpopulations based on transcriptional differences and visualized on Uniform Manifold Approximation and Projection (UMAP) plots, with the optimal number of clusters per samples determined using Clustree (Methods). Each tumour contained 3-8 transcriptionally distinct subpopulations of cells, with an average of 5 subpopulations per tumour [Sup Fig S2]. Functional enrichment analysis using the MSigDB Hallmarks and CancerSEA signatures (Yuan et al., 2019) revealed transcriptional subpopulations enriched for similar gene sets across all tumours, with proliferation and stemness signatures seen in at least one cluster in every tumour and EMT, hypoxia and invasion signatures represented as well [Sup Fig S2]. These enrichments may represent common biological processes active across all neuroendocrine pathologies in prostate cancer. In contrast, the degree of transcriptional heterogeneity varied with tumour pathology. While most PDXs had 2-3 distinct neuroendocrine clusters, all small cell NE PDXs had 5-6 subpopulations no matter whether they were derived from primary (224R and 224R-Cx) or metastatic (435.31A-Cx) tissues.

Pathology also determined the clustering of focal NED and mixed adeno-small cell tumours, with cells with adenocarcinoma markers forming distinct clusters from cells expressing neuroendocrine markers. In the focal NED PDXs 272R and 470B, neuroendocrine clusters were located on the UMAPs in close proximity to adenocarcinoma clusters, but in the mixed adeno-small cell PDX 224R, these two populations were clearly distant from one another [Sup Fig S2]. Such differences in clustering patterns with individual PDXs suggest each neuroendocrine pathology may harbour its own set of transcriptionally distinct sub-populations.

### Distinctive transcriptional subpopulations distinguish different neuroendocrine pathologies from each other

To identify transcriptional sub-populations shared between or unique to NE pathologies, we adopted a data integration strategy based on the CSS Simspec method, which showed optimal ability to match cells based on pathology in our benchmarking studies using the 224R and 224R-Cx samples [Methods; Supplementary Note 1]. Integrating expression counts from 18,632 cells from all 9 PDXs using CSS Simspec yielded 16 clusters. The positions of these clusters on the UMAP reflected differences in tumour pathology (Fig 2A), which had greater influence on clustering than cell cycle state, prior treatment status or site of tissue collection [Sup Fig S3].

**Figure 2.**
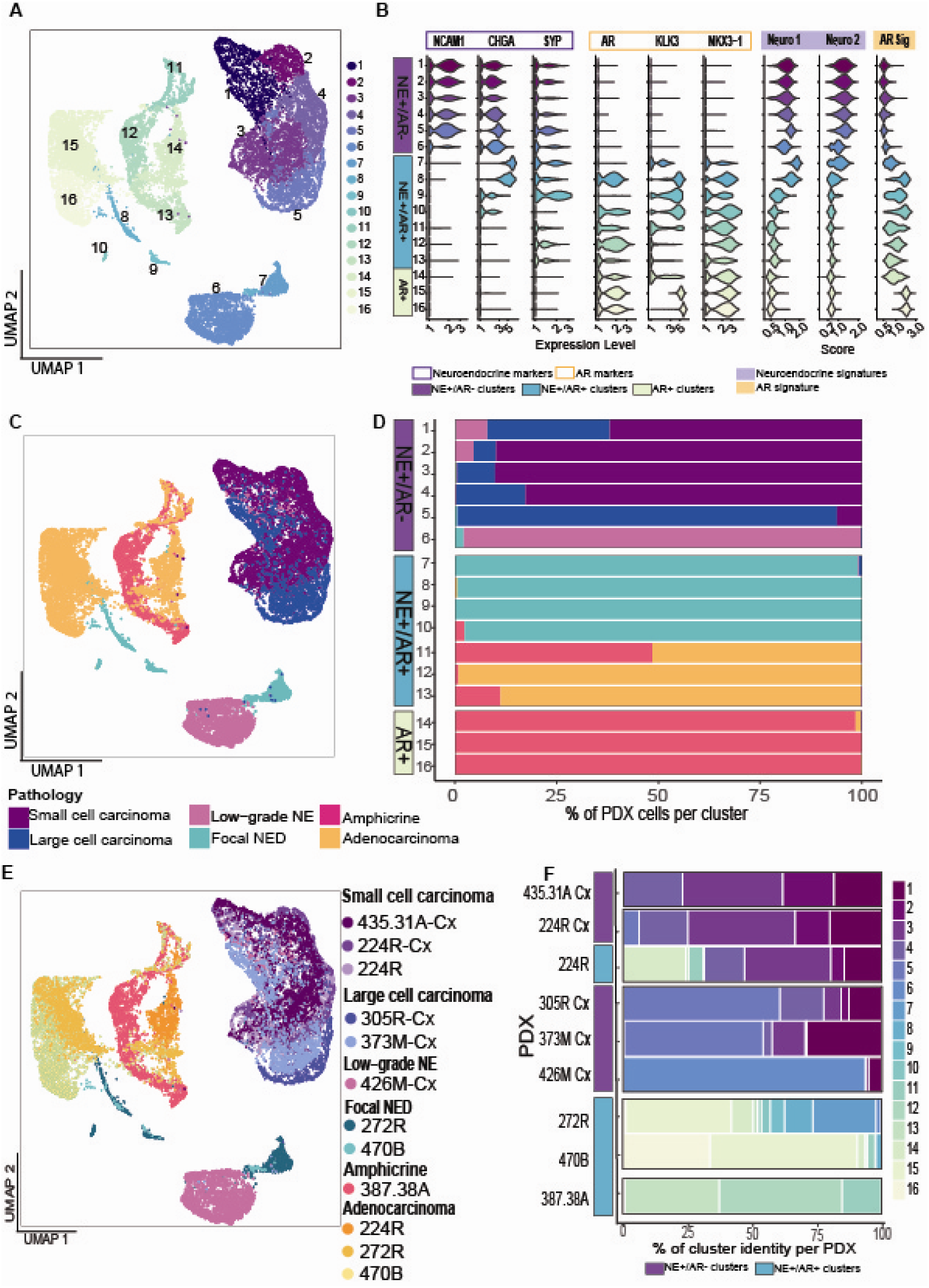
Inter-tumoural heterogeneity can be observed between the different pathologies and patients. A) UMAP depicting the multiple sub-clusters detected in the integrated dataset. 16 clusters were detected. B) Violin plot showing the range of expression of the neuroendocrine-specific genes (SYP, CHGA, NCAM1) and androgen-regulated genes (AR, KLK3, NKX3.1) per cluster. Clusters 1-6 are labelled as NE+/AR-, clusters 7-13 are labelled as NE+/AR+ and clusters 14-16 are AR+/NE-. C) UMAP shows the location of each sample. Clusters 1-6 comprise small and large cell pathologies. Clusters 7-13 include mixed pathologies (NED and amphicrine). Clusters 14-16 consists in adenocarcinoma cells. D) Stacked bar plot representing the contribution of each tumour to the individual clusters. E) UMAP coloured by PDX sample. F) Stacked bar plot describing the proportion of clusters per PDX sample.

Clusters 1-6 displayed robust expression of the neuroendocrine markers NCAM1, CHGA and SYP and virtually no expression of AR signalling markers (Fig 2B): thus, they were labelled NE+/AR- clusters. Cells in Clusters 1-6 were predominantly from tumours with large or small cell NE pathologies (Figs 2C & 2D). Clusters 1-4 were shared across PDXs with small cell pathology (224R, 224R-Cx and 435.31A-Cx) and large cell pathology (305R-Cx and 373M-Cx) (Figs 2E & 2F), revealing an overlap in the composition of some transcriptional sub-populations between these types of NEPC. In contrast, cluster 5 predominantly contained cells from large-cell NEPC tumours (305R- Cx and 373M-Cx). Nearly all the cells in cluster 6 were from PDX 426M-Cx, which has low-grade NE pathology. This cluster was situated far from the other NE+/AR- populations, likely reflecting the unique clinical characteristics of the patient. Clusters 1-5 contained a high proportion of cells in S and G2M phase, reflecting the highly proliferative nature of fully differentiated NEPC. [Sup Fig S3A].

Clusters 7 – 13 co-expressed neuroendocrine and AR signalling markers (NE+/AR+; Fig 2B). Each of these clusters displayed variable expression of CHGA and/or SYP, but little to no NCAM1. Similarly, expression of AR and KLK3 varied across these clusters. Most cells in NE+/AR+ clusters came from PDXs with the intermediate focal NED and amphicrine pathologies (Figs 2C & 2D), revealing focal NED to also be an AR-expressing neuroendocrine pathology. Clusters 8-10 were specific to tumours with focal NED pathology (PDX 272 and 470B) while clusters 11-13 were from the amphicrine tumour (Fig 2E & 2F) indicating that although they share AR expression, focal NED and amphicrine pathologies diverge at the transcriptional level from each other.

Finally, Clusters 14-16 had robust expression of AR-related genes with virtually no NE gene expression (AR+; Fig 2B). They were comprised of cells from the adenocarcinoma component of the focal NED tumours and the mixed adeno-small cell PDX 224R.

### Cells with focal NED pathology co-express neuroendocrine and adenocarcinoma markers

Detection of both AR signalling and neuroendocrine genes in clusters 8-10 could be linked to presence of cells with concurrent expression of both sets of genes, but could also occur if those clusters contained a mix of neuroendocrine and adenocarcinoma cells. To investigate, we performed cell-level co-expression analysis using the “Featureplot” function of Seurat to enumerate and visualize the fraction of cells in a sample with detectable expressed of both markers AR signalling genes and canonical NE markers within individual neuroendocrine cells. PDX 272R, which has focal NED pathology, was analysed along with the amphicrine PDX 387.38A and the mixed adeno-small cell PDX 224R as positive and negative controls for co-expression of AR signalling and NE genes, respectively.

As expected of an amphicrine tumour, PDX387.38A displayed strong co-expression of *SYP* with multiple adenocarcinoma markers (Figure 3A). Transcript counts for each pair of markers was generally robust in the cells where co-expression was detected. In contrast, PDX 224R with the mixed of small cell and adenocarcinoma pathology displayed very limited co-expression of its most abundant NE marker gene, *ASCL1*, and AR signalling genes (Fig 3B), in line with separation of its adenocarcinoma and neuroendocrine components on a UMAP plot [Supp Fig S2] PDX 272R displayed robust expression of multiple neuroendocrine as well as adenocarcinoma markers [Sup Fig S4], with *CHGA* being the neuroendocrine gene with highest average level of expression. *AR* and *CHGA* were concurrently expressed by 24.5% of cells in PDX 272R, while *KLK3-CHGA* co-expression was found in 60% of cells and *NKX3.1-CHGA* co-expression in 49.1% of cells (Fig 3C). Rates of co-expression of *CHGA* with AR markers in 272R exceeded those of the amphicrine PDX 387.38A, demonstrating that the focal NED component of 272R expressed the AR.

**Figure 3.**
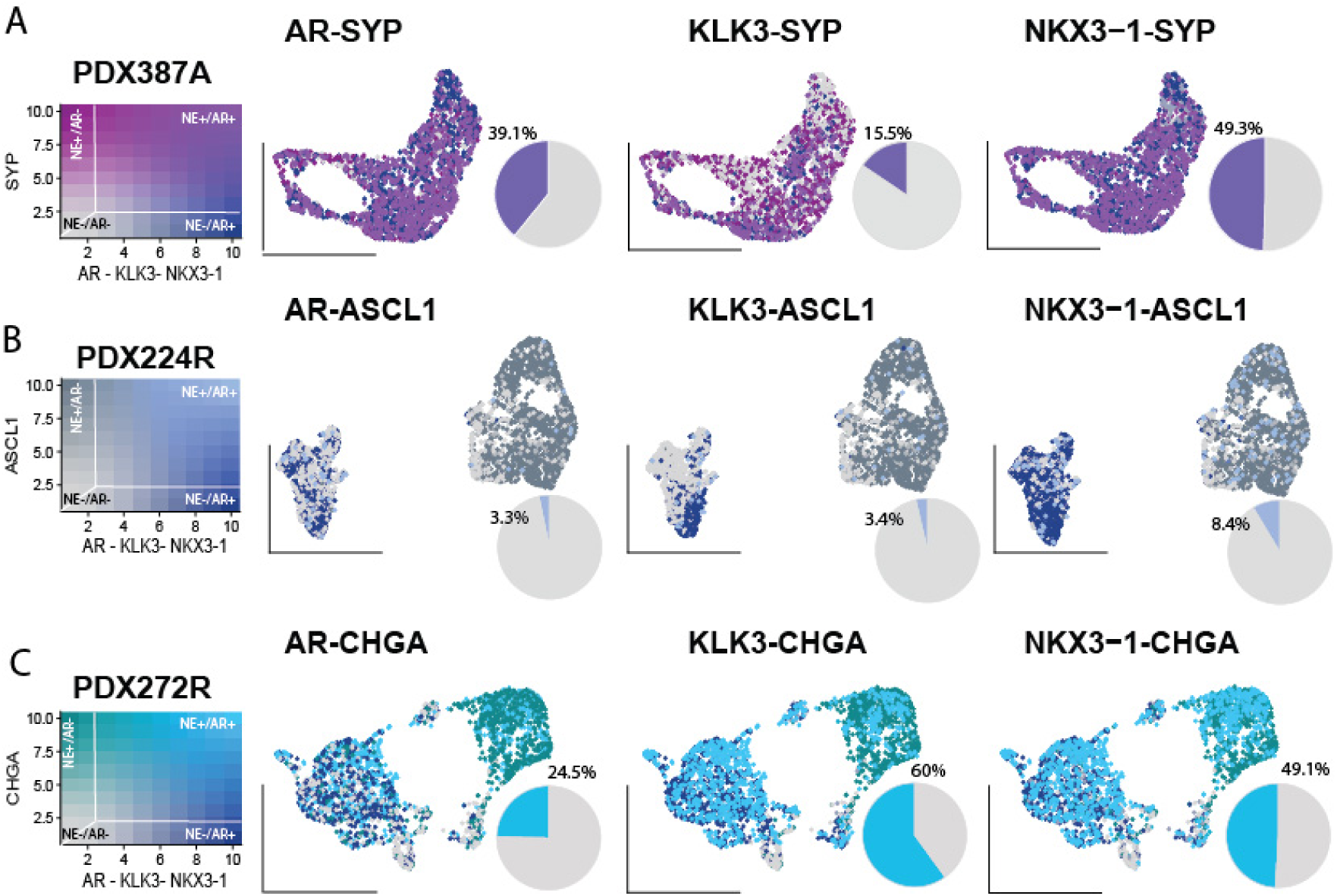
Co-expression analysis of neuroendocrine and adenocarcinoma markers. Colour blending represents the co-expression level; data has been scaled from 0-10. Zero represents cells without any expression of the markers, while 10 represents cells with the highest expression level. The percentage represents only the cells that co-express such markers. **A)** UMAPs representing the cells that co-express AR-SYP, KLK3-SYP, and NKX3.1-SYP in sample PDX387.38A. Pink represents cells expressing SYP, and dark blue represents cells expressing AR, KLK3 or NKX3.1. A purple shade represents cells that highly co-express such markers. Grey shades represent cells that don’t express any marker. **B**) UMAPs representing the cells that co-express AR-CHGA, KLK3- CHGA, and NKX3.1-CHGA in sample PDX272R. Green colour represents cells that uniquely express CHGA, and dark blue represents cells that uniquely express AR or KLK3 or NKX3.1. A blue shade represents cells that highly co-express such markers. Grey shades represent cells that don’t express any marker. **C**) UMAPs representing the cells that co-express AR-ASCL1, KLK3-ASCL1, and NKX3.1-ASCL1 in sample PDX224R. Dark grey represents cells that uniquely express ASCL1, and dark blue represents cells that uniquely express AR or KLK3 or NKX3.1. A light blue shade represents cells that highly co-express such markers. Grey shades represent cells that don’t express any marker.

Interestingly, co-expression of *CHGA* and *AR* regulated genes was also observed in the adenocarcinoma component of PDX 272R. However, cells from PDX 287R, a pure adenocarcinoma from the MURAL collection profiled in a recent study (Porter et al, 2023), displayed virtually no detectable expression of NE genes [Supp Fig S5]. Co-expression of both markers in prostate adenocarcinoma may therefore only occur in the context of focal neurodifferentiation.

### Unique expression signatures distinguish types of neuroendocrine pathologies in prostate cancer

To compare transcriptional differences between neuroendocrine pathologies, we applied differential gene expression and gene set enrichment (GSE) analysis to all 16 clusters in the integrated data set. There was high overlap in the top 5 differentially expressed genes (DEGs) within the NE+/AR- clusters, which represent the small and large-cell NE pathologies that lack AR expression (Fig 4A). Very few DEGs were shared with the NE+/AR+ clusters, which contain the AR-expressing focal NED and amphicrine populations. The exceptions were cell cycle genes such as *MKI67*, which overlapped between Clusters 1, 2 and 11 due their proliferative nature. Comparison of DEGs suggested AR-expressing neuroendocrine pathologies have distinct transcriptional signatures from those that lack AR expression.

**Figure 4.**
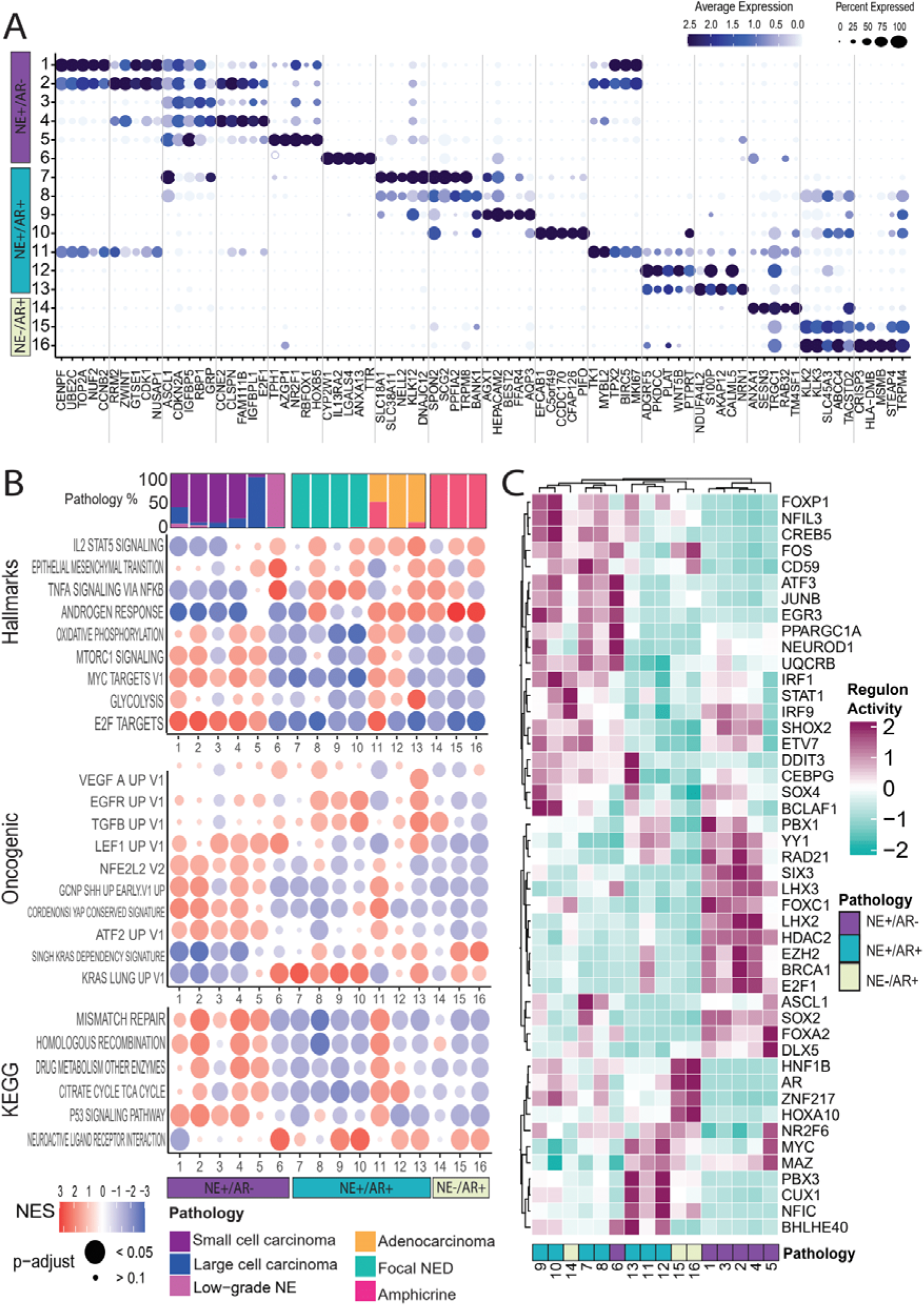
Characterizing gene expression features that distinguish neuroendocrine sub- populations within the integrated data set. A) Genes that are differentially expressed per cluster are shown. The size of the circle represents the percentage of cells expressing the gene. The intensity of the colour represents the gene’s expression level; dark blue signifies a higher level, while light blue to white shows a low or null expression of the gene. **B)** Enrichment of selected Hallmark, Oncogenic targets and KEGG gene sets from MSigDB are shown. The size of the circle represents adjusted p-value, small circles represent p-values over 0.1, and larger circles represent p-values less than 0.05. The enrichment score is represented by colour; red is a positive enrichment score, and blue is a negative enrichment score. **C)** Transcription factor analysis heatmap. Regulon activity is scored from 2 to -2, where 2 represents a positive regulon activity, and -2 symbolises a negative regulon activity. Here 46 manually curated TFs out of the 87 significantly enriched TFs reported by SCENIC are displayed.

To search for differences in cancer-related pathways and processes between neuroendocrine pathologies, cluster-level enrichment was assessed for the Hallmarks 50, oncogenic and Kyoto Encyclopedia of Genes and Genomes (KEGG) gene sets from MSigDB (Liberzon et al., 2015). Cells from AR+/NE- small and large cell neuroendocrine pathologies have distinct enrichment profiles from AR+/NE+ amphicrine and focal NED cells (Fig 4B). Certain gene sets showed mutually exclusive patterns of enrichments in focal NED as compared to small and large-cell pathologies, while clusters from the amphicrine PDX shared gene set signatures with both focal NED and small and large-cell PDXs.

The common set of enriched gene sets was observed in the NE+/AR- Clusters 1-5 included E2F targets and oxidative phosphorylation, consistent with the highly proliferative nature of small and large cell NEPC. Noteworthy oncogenic signalling enrichments included MYC, MTORC1, LEF1, a key regulator of epithelial-mesenchymal transition (EMT), and YAP signalling, which was recently linked to emergence of stemness phenotypes in castration-resistant prostate cancer (Tang et al, 2022).

In contrast, the NE+/AR+ Clusters 7-10 that are dominated by the focal NED pathology displayed markedly different enrichment from Clusters 1-5. Top enrichments included the TNFA signalling via NFKB and KRAS signalling, both of which were also enriched in the adenocarcinoma Clusters 14-16 but downregulated in the small/large cell clusters (Fig 3B). In contrast to adenocarcinoma, there was high expression of genes upregulated by EGFR and TGFβ. Enrichment of EMT was seen as well, though in focal NED activation of EMT may occur through KRAS instead (Kim et al., 2015). Indeed, each pathology shows divergent expression of EMT genes [Supp Fig S6]. Androgen response genes were most strongly upregulated in Cluster 8, indicating AR signalling expression may vary across focal NED cells.

Interestingly, Clusters 11-13 primarily represent cells from the amphicrine PDX 387.38A showed enrichment signatures in common with both Clusters 1-5, which include the NE+/AR- small and large cell NE pathologies, and Clusters 7-10, which were mainly NE+/AR+ focal NED (Fig 4B). Cluster 11 shared a GSE profile with Clusters 1-5, Cluster 12 matched all the other neuroendocrine pathologies, while Cluster 13 was most similar to focal NED as well as adenocarcinoma cells (Clusters 14-16). Thus, the amphicrine PDX 387.38A contains a mix of transcriptional states covering both AR-expressing pathologies as well as neuroendocrine pathologies where AR expression is suppressed.

Cluster 6, comprised nearly entirely of cells from the low-grade NE PDX 426M-Cx, was an outlier in the GSE analysis. Its GSE profile was more similar to Clusters 7-10 than to Clusters 1-5, including enrichments for TNFA signalling via NFKB, EMT, KRAS signalling and KEGG neuroactive ligand- receptor interactions. Despite little to no AR expression, PDX 426M-Cx appears to share some characteristics with AR-expressing pathologies.

Transdifferentiation to neuroendocrine prostate cancer is driven by key master regulator transcriptional factors (TFs) (Mu et al., 2017; Adams et al. 2019; Guo et al., 2019). To infer how changes to gene regulatory networks contribute to differences in gene expression states observed between the 16 cell clusters in our data, we scored activity of transcription factor regulons in each cluster using the SCENIC algorithm. Concordant with prior results, the small/large cell clusters (1-5) and focal NED clusters (7-10) showed clear divergence in inferred TF regulon activity (Fig 4C). Clusters 1-5 were predicted to have high activity of several TFs that regulate proliferation, chromatin state and DNA replication/repair, including E2F1, EZH2, HDAC2 and BRCA1. Regulons for known neuroendocrine lineage regulators ASCL1, SOX2 and FOXA2 were active in all small cell, large cell and focal NED clusters, but scored markedly higher in the small/large cell clusters. There were numerous TFs with high activity in focal NED Clusters 7-10 but weak to no activity in the small/large cell clusters (Fig 4C). Among the TFs specific to focal NED clusters were the stemness factors FOS and JUNB, along with NEUROD1, a TF shown to contribute to global transcriptional differences between NEPC tumours (Labreque et al., 2010). The focal NED clusters also showed overlap in TF activity with the adenocarcinoma clusters (14-16), aligning with the observation of retained AR signalling in those cells. As expected, AR was one of the shared TFs but there were also others, including FOS and the lineage factor HOXA10. As in the GSE analysis, the amphicrine clusters 11- 13 showed overlap in TF activity with both small/large cell and focal NED clusters. ASCL1 was not active in amphicrine clusters, however. SCENIC found 6 TFs (DDIT3, CEBPG, PBX3, CUX1, NFIC and BHLHE40) with elevated activity in amphicrine clusters.

In summary in addition to AR signalling, several other biologically meaningful differences expression were seen between the AR-expressing focal NED and amphicrine pathologies and small and large cell neuroendocrine cells, which lack AR activity. These included numerous pathways and processes involved in oncogenic signalling, inflammation, and metastasis. Differences in TF activity between pathologies were predicted by SCENIC, with the focal NED and amphicrine pathologies showing overlap in TF regulon expression with adenocarcinoma cells.

### Single-cell copy-number profiling indicates focal NED and small cell neuroendocrine carcinoma arise by different means in tumours of mixed pathology

Transcriptional profiling of single-cell clusters supports the small cell neuroendocrine pathology as being further diverged from adenocarcinoma than the focal NED pathology. This is reflected in the single-sample UMAPs from PDXs 224R and 272R [Supp Figs S3 and S4]. As noted, the adenocarcinoma and neuroendocrine subpopulations occupy separate and distant regions of the UMAP plot for 224R, indicative of divergent cell states. However, in the UMAP 272R the adenocarcinoma and focal NED cells cluster close together in a nearly contiguous mass consistent with a continuum of cell states. This view is further supported by pseudotime analysis [Supp Fig S7] as well as the observed co-expression of AR and NE markers in cells of 272R but not 224R [Fig 3].

To determine whether genetic divergence existed between the adenocarcinoma and neuroendocrine subpopulations in PDXs 224R and 272R, we inferred copy-number status at the chromosome arm level from the transcriptomes of individual cells. Following the methods of Kinker et al we searched for genetically distinct clonal subpopulations within each PDX tumour. Briefly, combined expression of genes on the same chromosome arm is measured in each cell to detect heterogeneity in expression level chromosome arms indicative of copy-number gains and losses in a sample. Clustering cells by arm-level expression detects clonal subpopulations with genetic differences at the copy-number level [Methods].

Three distinct clonal sub-populations could be detected in PDX 224R on the basis of inferred copy-number states on four chromosome arms: 21q, 8q and 9p and 7q [Fig 5A]. Clones 1 and 3 mapped exclusively to the neuroendocrine cell clusters of 224R, while Clone 2 was found only in the adenocarcinoma clusters [Fig 5B and 5D]. Expression of genes on 21q, 8q, 7q and 9p showed consistent differences across all neuroendocrine and adenocarcinoma clusters [Fig 5C], indicating the neuroendocrine and adenocarcinoma cells in PDX 224R come from genetically distinct clones.

**Figure 5:**
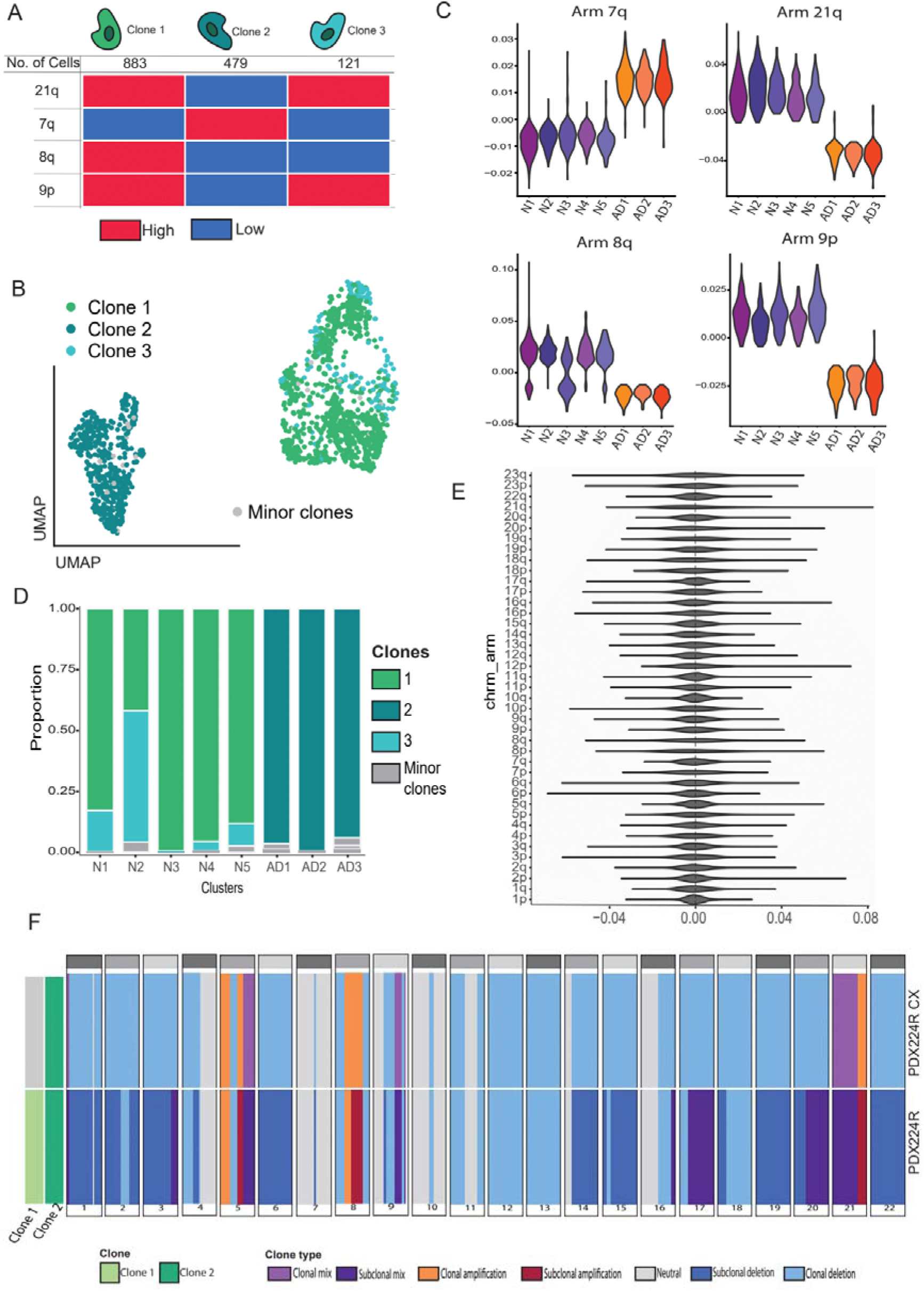
The small cell neuroendocrine and adenocarcinoma components of the mixed pathology in PDX224R comprise genetically distinct clonal sub-populations. (**A**) Copy-number profiling of cells based on combined expression levels of genes per chromosome arm from single-cell RNA-sequencing identifies three genetically distinct clonal sub-populations distinguished by differences on four chromosomal arms. (**B**) UMAP coloured by clone. Adenocarcinoma clusters are represented mainly by clone 2, while neuroendocrine clusters are represented by clone 1 and 3. (**C-D**) The neuroendocrine and adenocarcinoma components of PDX224R differ in expression level of genes on all four of the chromosome arms that define clones. Clones 1 and 3 are found exclusively in the neuroendocrine sub-populations while Clone 2 exclusively belongs to the adenocarcinoma sub- populations. (**E**) No evidence could be found for existence of distinct genetic clones within the focal NED PDX272. (**F**) Whole-genome sequencing of PDX224R and PDX224R-Cx reveals loss of clonal diversity after castration and retention of a clone matching the profile of the genetic clone overlapping the neuroendocrine population within PDX224R single-cell RNA-sequencing data.

In contrast, cells from PDX 224-Cx, which was derived from 224R via growth in castrate host mice, showed no consistent or substantial differences in gene expression at the chromosome arm-level [Fig 5E], suggestive of selection and emergence of a single clone from the neuroendocrine population post-castration. To validate these findings, whole-genome sequencing (WGS) to 80-90X coverage [Methods] of 224R, 224R-Cx and a germline sample from patient 224 was performed and analysed using the HATCHet algorithm, which infers clonal subpopulations from WGS based on frequencies of copy-number alterations across matched samples from the same patient (Zaccaria and Raphael, 2020). HATCHet predicted the presence of two distinct clonal sub-populations in 224R differing in copy-number profile across 17 of the 22 autosomes, including copy-number changes on 7q, 8q, 9p and 21p [Fig 5F]. In contrast, only a single clone was detected in PDX 224R-Cx. The patterns of amplification and deletion on 7q, 8q, 9p and 21p were concordant with those inferred from single-cell RNA-sequencing data.

In contrast, no differences in expression at the chromosome arm level were detected from the 272R single-cell RNA-sequencing data, indicating this PDX is homogeneous at copy-number level, harbouring only a single dominant clone [Fig 5E] Therefore, the adenocarcinoma and focal NED populations in 272R come from the same clone. The presence of a cluster with low expression of both AR and NE markers between the adenocarcinoma and focal NED clusters marks 272R as actively undergoing a process of transdifferentiation [Supp Fig S3]. PDX 470B also has focal NED pathology and likewise showed no evidence for copy-number differences amongst adenocarcinoma and focal NED cells [Supp Fig S8]. On the other hand, the genetic differences between the adenocarcinoma and small cell neuroendocrine subpopulations in PDX 224R are more consistent with divergence of two populations in the prostate prior to diagnosis. These contrasting patterns of sub- clonality align with the concept of focal NED and small cell neuroendocrine pathologies as being distinct entities within a spectrum of neuroendocrine states, with focal NED being less diverged from adenocarcinoma.

## Discussion

Integrative analysis of the transcriptional profiles of 18,632 individual cells from nine PDXs of NEPC demonstrated transcriptional features of neuroendocrine cells are strongly associated with pathology. Strikingly, focal NED cells retain expression of AR signalling genes at levels comparable to amphicrine and adenocarcinoma cells, while maintaining robust expression of neuroendocrine markers (Figure 2). Co-expression of the neuroendocrine marker CHGA along with one of more of AR, KLK3 and NKX3.1 was widespread in focal NED cells (Figure 3). Thus, like amphicrine, focal NED is another neuroendocrine pathology with capacity for AR signalling.

Neuroendocrine pathologies that retain AR signalling have distinct patterns of intra-tumoural transcriptional heterogeneity from those do not, involving multiple oncogenic processes. The AR-null small and large cell NE pathologies displayed marked upregulation of growth-associated processes such as Myc and YAP signalling, DNA repair and oxidative phosphorylation relative to other pathologies (Figure 4B), consistent with their more proliferative nature. In contrast, these signatures were depleted in focal NED, amphicrine and the low-grade NE sub-populations, which instead were enriched for a non-overlapping set of pathways, including KRAS, TNF-alpha, EGFR and IL2-STAT5 signalling. Similarly, each pathology showed a unique profile of activity of master regulator transcription factors (Figure 4C).

Transcriptional sub-populations expressing EMT genes were observed in every PDX regardless of pathology [Figure 4, Supp Figs S2 and S6]. The well-established roles of EMT in plasticity and metastasis may underlie the aggressiveness of NEPC. Notably, the activation of EMT exhibits pathology-specific patterns. Focal NED cells were enriched for KRAS signaling, recognized as an EMT driver (Brabletz et al., 2018; Thiery et al., 2009), while large and small cell pathologies prominently express LEF1, another acknowledged EMT activator (Liang et al., 2015).

Whether focal NED can transition to small or large cell pathology remains unresolved. The focal NED PDXs 272R and 470B contained intermediate transcriptional sub-populations between adenocarcinoma and focal NED. In contrast, within the mixed small cell-adenocarcinoma PDX 224R the neuroendocrine and adenocarcinoma sub-populations were both transcriptionally and genetically distinct (Figure 5), indicating complete trans-differentiation and long-standing divergence. Further longitudinal studies of PDXs and patients may shed light on whether focal NED is a transitional or terminally differentiated state.

Current standard of care chemotherapies for prostate cancer patients with neuroendocrine pathologies do not confer lasting benefit. Our results suggest new therapeutic options based on pathology. Focal NED may retain sensitivity to androgen-targeting agents and could respond to disruption of KRAS and EGFR signalling. In contrast, YAP and Wnt targeting agents may work better against small and large-cell NEPC tumours. Drugs targeting each of these pathways have shown effectiveness in solid tumours (Gibault et al. 2017; Liu et al., 2017; Mustachio et al., 2021; Tang et al. 2022; Zhang et al, 2020) but have not yet been deeply explored as therapeutic options for prostate cancer.

The MURAL PDX collection afforded an opportunity to isolate cells of rare NE pathologies and study them comprehensively at the transcriptional level. Our PDX models faithfully recapitulate the molecular profiles of the original donor tumours (Risbridger et al. 2021) and features of transcriptional ITH observed here are consistent with single-cell studies of CRPC (Bolis et al., 2021; Brady et al., 2021; Conteduca et al., 2021; Dong et al., 2020; Horning et al., 2018; Wang et al., 2022) and small-cell lung cancer (Stewart et al. 2020).

## Conclusions

Single-cell RNA-sequencing of a diverse spectrum of PDX models of NEPC reveals focal NED as being transcriptionally distinct from small and large cell NEPC, requiring its own treatment and management strategies. Our work redefines the molecular landscape in NEPC, revealing previously hidden layers of transcriptional heterogeneity that provide a basis to further develop new therapeutic opportunities for this low-survival subtype of prostate cancer.

## Declarations

### Ethics Statement

Patient-derived xenografts were established by the Melbourne Urological Research Alliance (MURAL) with informed written consent according to human ethics approvals from Monash Health (RES-20-0000-107C), the Peter MacCallum Cancer Centre (18/76; 11/102; 27/97), Eastern Health (E55/1213) and Monash University (12287). All animal handling and procedures were approved by the Monash University Standing Committee of Ethics in Animal Experimentation (MARP 2012/158, MARP/2014/085, MARP/2018/087 and 28911)

### Consent for publication

Not applicable.

### Availability of data and materials

R and Python code used to analyse data and generate figures included in this manuscript is available at https://github.com/dlgoode/QuezadaUrban_scRNAseq. The datasets generated and/or analysed during the current study are not yet publicly available due establishment of a controlled-access repository still being in progress, but transcript counts and processed data are available from the corresponding author on reasonable request.

### Authors’ contributions

RQU designed the work, performed analysis, interpreted data, prepared figures and wrote and revised the manuscript. SK performed analysis and prepared figures. AC generated data and tissue samples from xenografts and prepared figures. HW generated data and tissue samples from xenografts. BP advised on methodology. AB performed analysis. AR reviewed and assigned tumour pathology. HT contributed tissues for xenografts. RAT interpreted data, supervised the work and revised the manuscript. MGL interpreted data, supervised the work and revised the manuscript. GR interpreted data, supervised the work and revised the manuscript. RT conceived the study, designed and supervised the work, generated and interpreted data, prepared figures and revised the manuscript. DLG conceived the study, designed and supervised the work, interpreted data, performed analysis and wrote and revised the manuscript. All authors read and approved the final manuscript.

## Acknowledgements

We acknowledge the members of the Prostate Cancer Research program, the patients, families, and consumers who support our research. We acknowledge the patient representatives, clinical co- ordinators, scientists, and clinicians, who contribute to the Melbourne Urological Research Alliance (MURAL) and its collection of patient-derived models; the CASCADE program; and Kathleen Cuningham Foundation Consortium (kConFab) including kConFab research nurses and staff, the heads and staff of the Family Cancer Clinics, and the Clinical Follow-Up Study (which has received funding from the NHMRC, the National Breast Cancer Foundation, Cancer Australia, and the National Institute of Health (USA)) for their contributions to this resource, and the many families who contribute to kConFab. Patient derived xenograft models were established and maintained through the Melbourne Urological Research Alliance (MURAL). Library preparation and sequencing were provided by the Single Cell Innovation Laboratory, Centre for Cancer Research, University of Melbourne and the Molecular Genomics Core Facility, Peter MacCallum Cancer Centre. This research was supported by the Monash University Histology Platform, Monash University Animal Research Laboratories, the Research Computing Facility at the Peter MacCallum Cancer Centre and the NeCTAR Research Cloud.

