## Supplemental material for "Defining focal neuroendocrine differentiation as a transcriptionally distinct form of prostate cancer pathology characterized by the expression of androgen receptors"

**This PDF file includes:**

Figures S1 to S12

Supplementary Note 1

**Supplementary Figures**

**A**

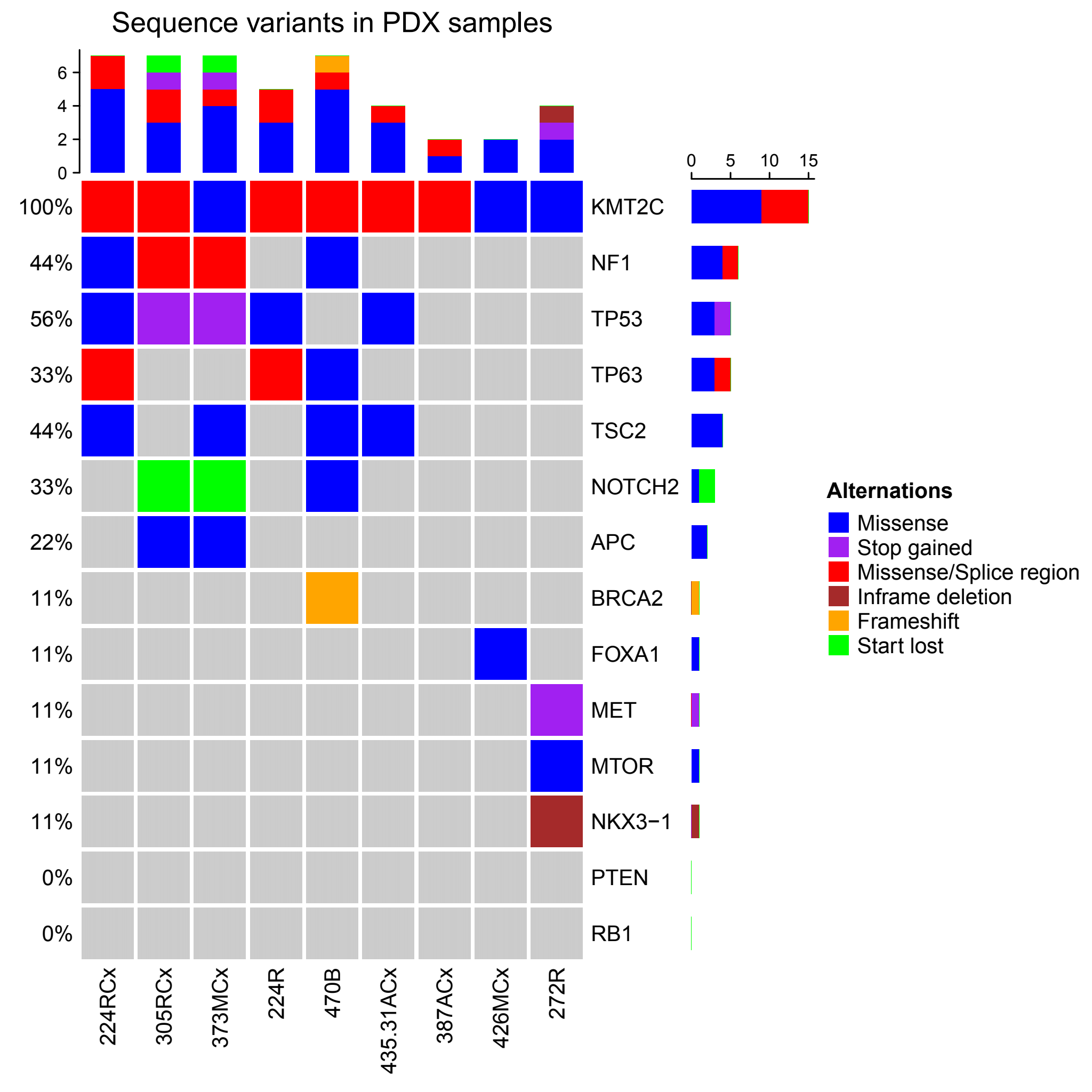

**B**

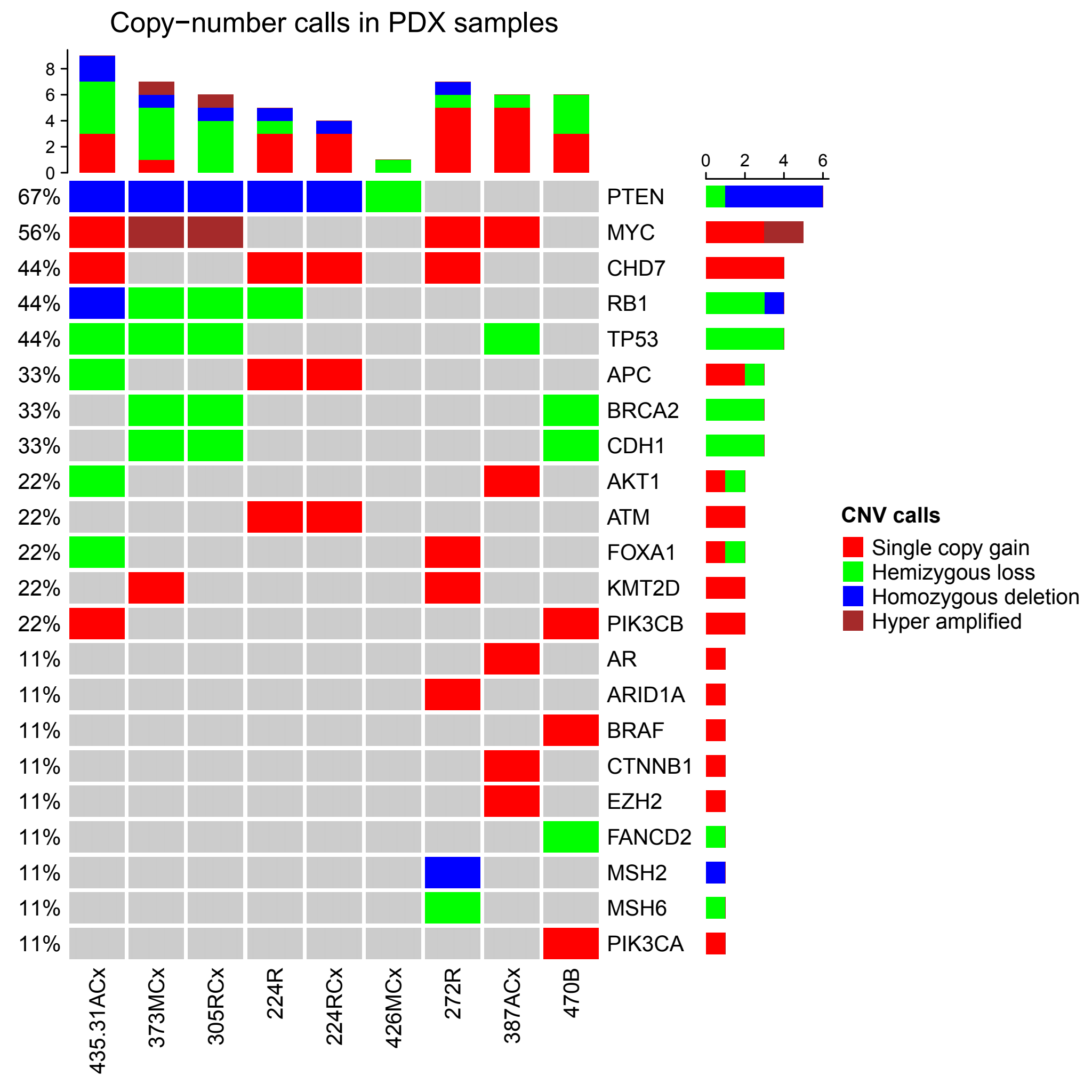

Figure S1 – Mutational profiles of the patient-derived xenograft models used in this study. (A) Protein-coding mutations in selected cancer driver genes for each PDX model, coloured by effect on protein sequence. Sidebar displays overall frequency of mutations to each gene across the 9 PDXs. (B) Copy-number aberrations in selected cancer driver genes for each PDX model coloured by magnitude of change relative to diploid.

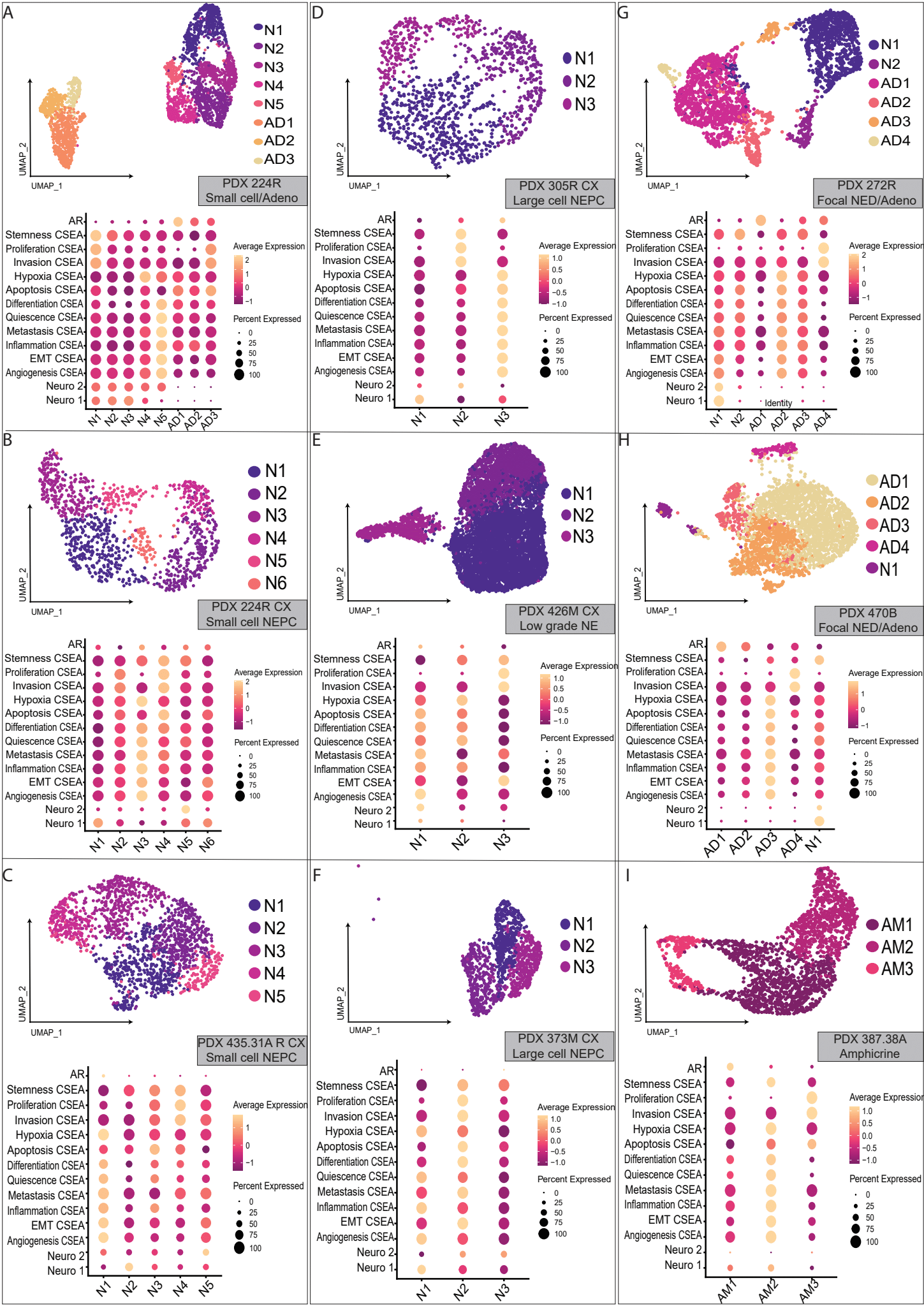

Figure S2 – UMAPs displaying cell clusters identified in each PDX, with corresponding gene set enrichment scores for functional states from the Cancer Single-cell Expression Atlas (CSEA) (Yuan et al, 2018) and androgen receptor (AR) signaling and neuroendocrine prostate cancer (Neuro1 and Neuro2) signatures from Labrecque et al, 2019. Clusters are labelled according to whether cells express neuroendocrine markers (N), adenocarcinoma markers (AD) or both sets of markers (AM). PDXs are divided according to pathology. (A) mixed small cell-adenocarcinoma PDX; (B) Small cell neuroendocrine prostate cancer (NEPC) PDX 224R-Cx; (C) Small cell NEPC PDX 435.31A-Cx; (D) Large cell NEPC PDX 305-Cx; (E) Low-grade neuroendocrine PDX 426M-Cx; (F) Large cell NEPC PDX 373M-Cx; (G) Adenocarcinoma with neuroendocrine differentiation (Adeno-NED) PDX 272R; (H) Adeno-NED PDX 470B; (I) Amphicrine PDX 387.38A.

**A**

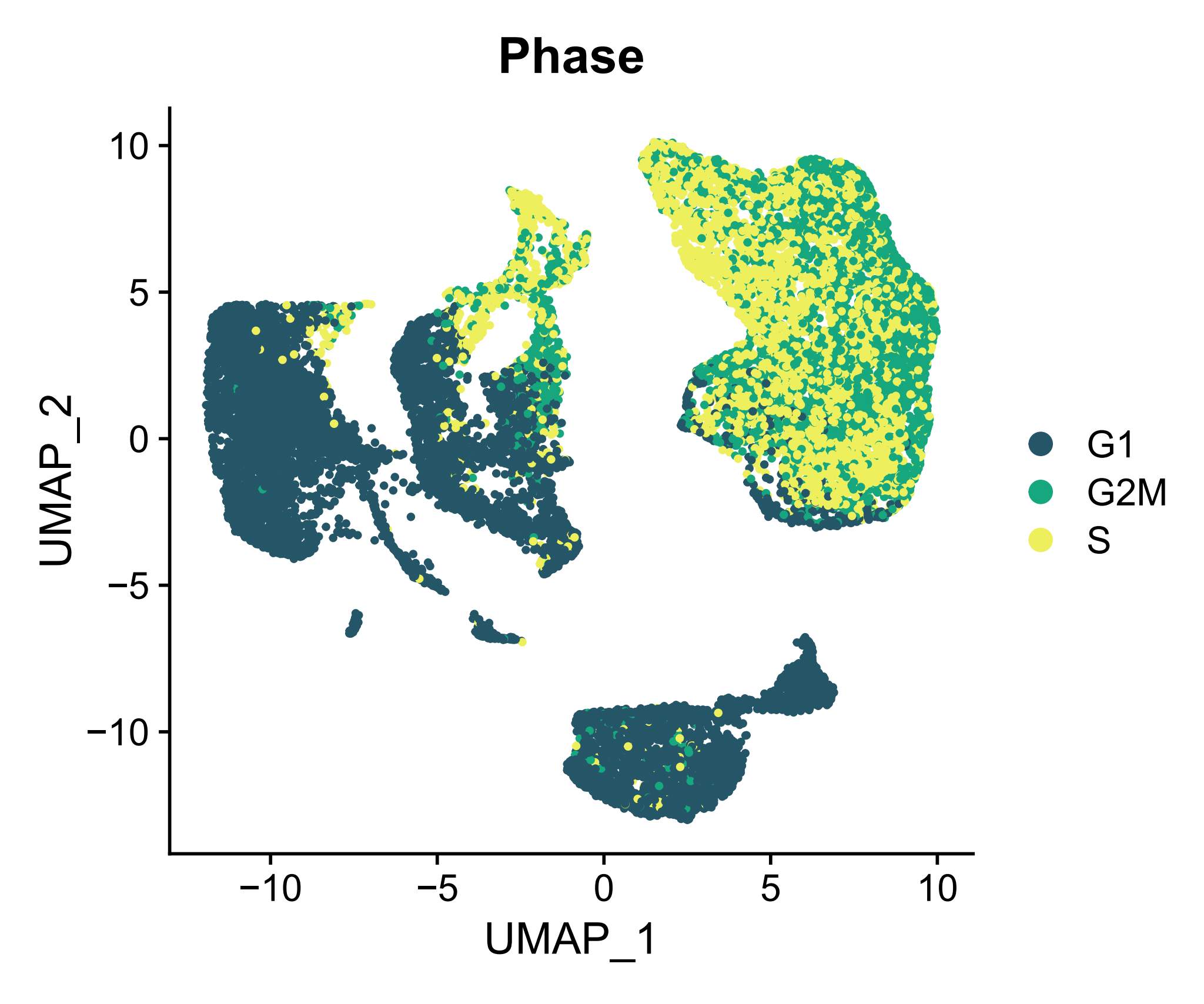

**B**

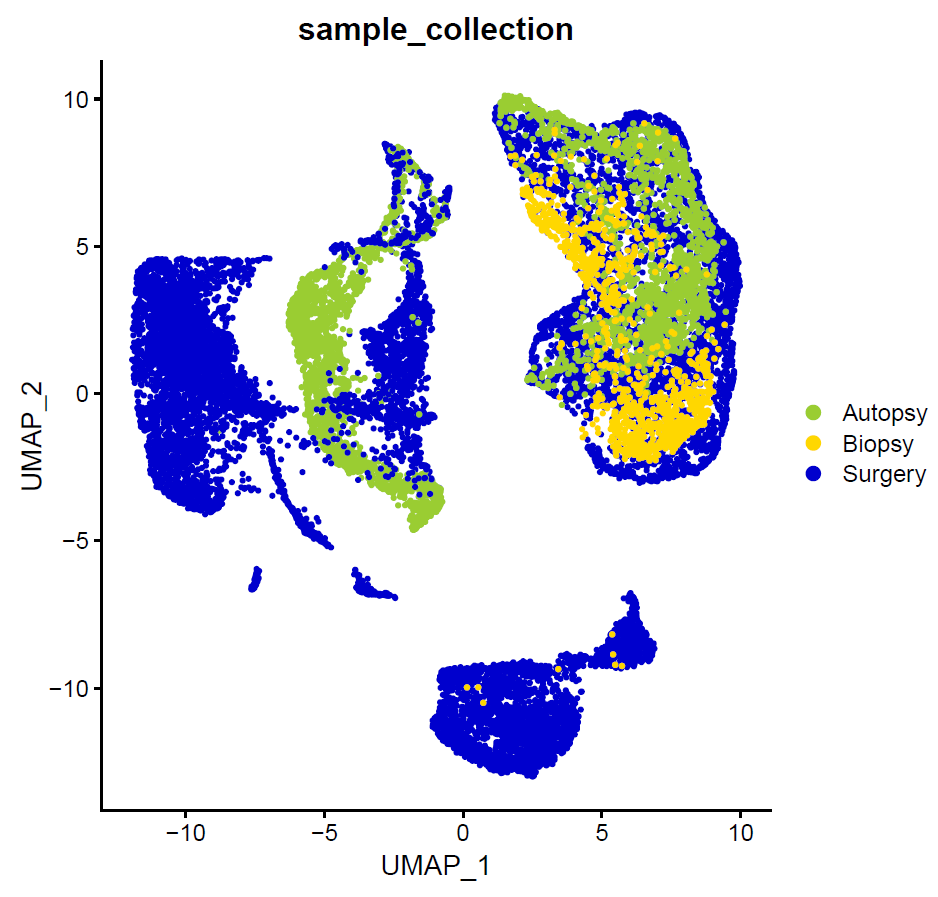

**C**

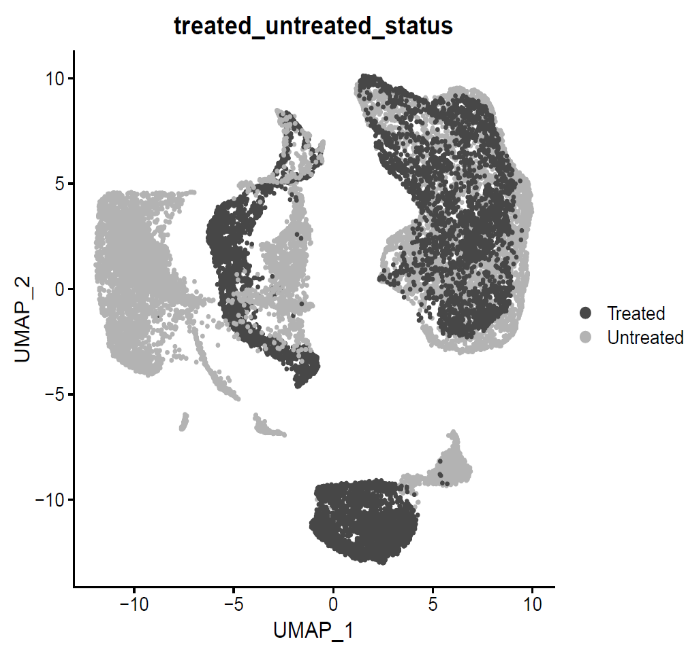

Figure S3 – UMAPs of integrated data from all 9 PDXs with cells coloured by (A) cell cycle phase, (B) site of collection or (C) treatment status.

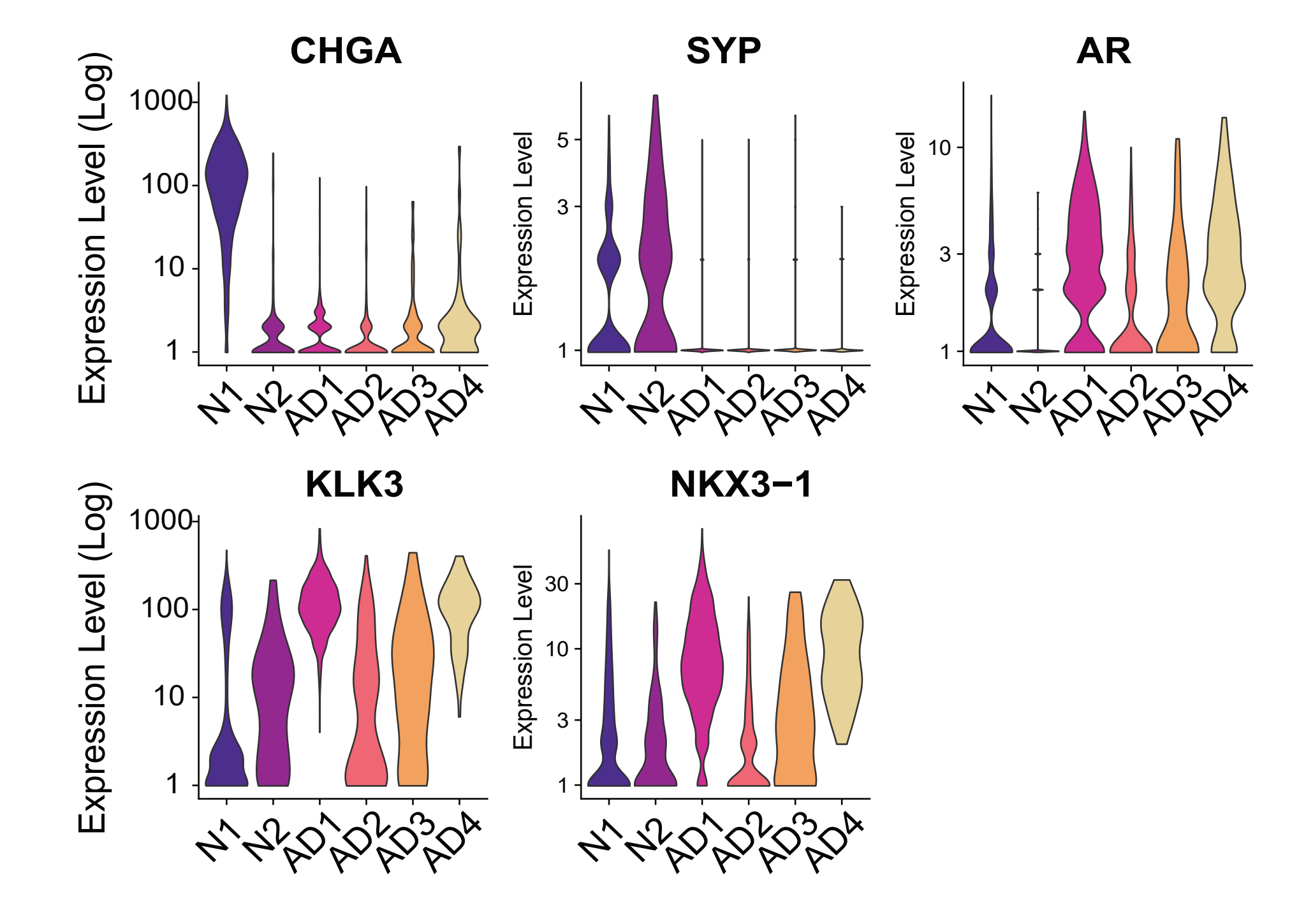

Figure S4 – Expression of neuroendocrine markers genes and CHGA and AR, and the adenocarcinoma marker genes by UMAP cluster in PDX 272R.

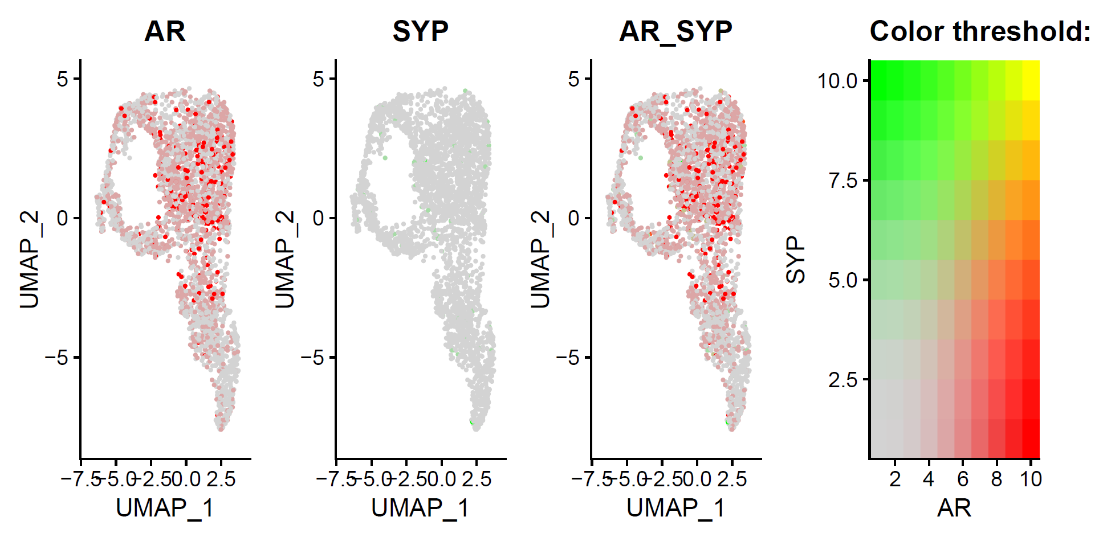

Figure S5 – Expression of the androgen receptor (AR) and synaptophysin (SYP) genes in tumour cells from PDX 287R.

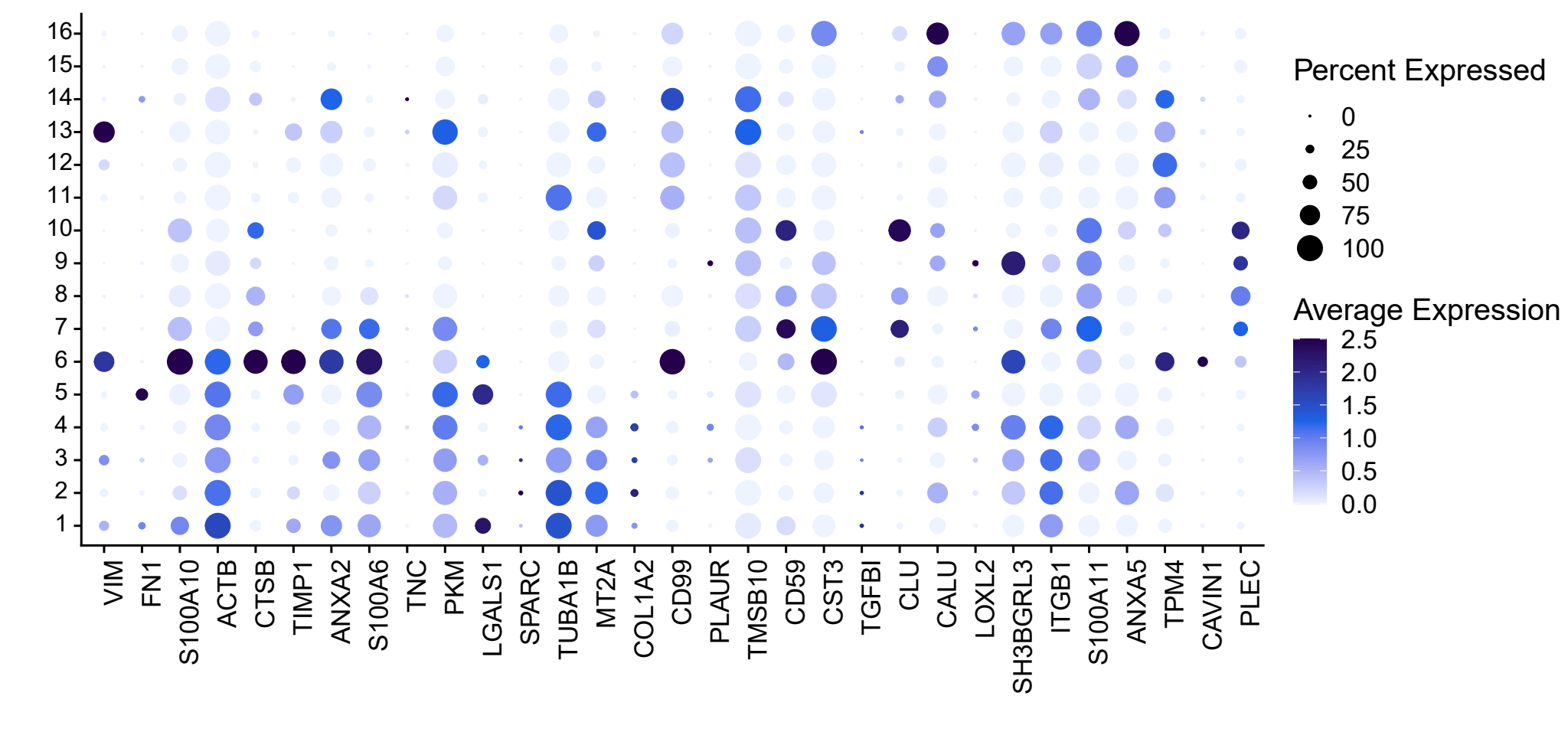

Figure S6 – Expression of genes from the Epithelial Mesenchymal Transition gene set from the Cancer Single-cell Expression Atlas (Cancer SEA) across UMAP clusters from the integrated analysis of PDX data.

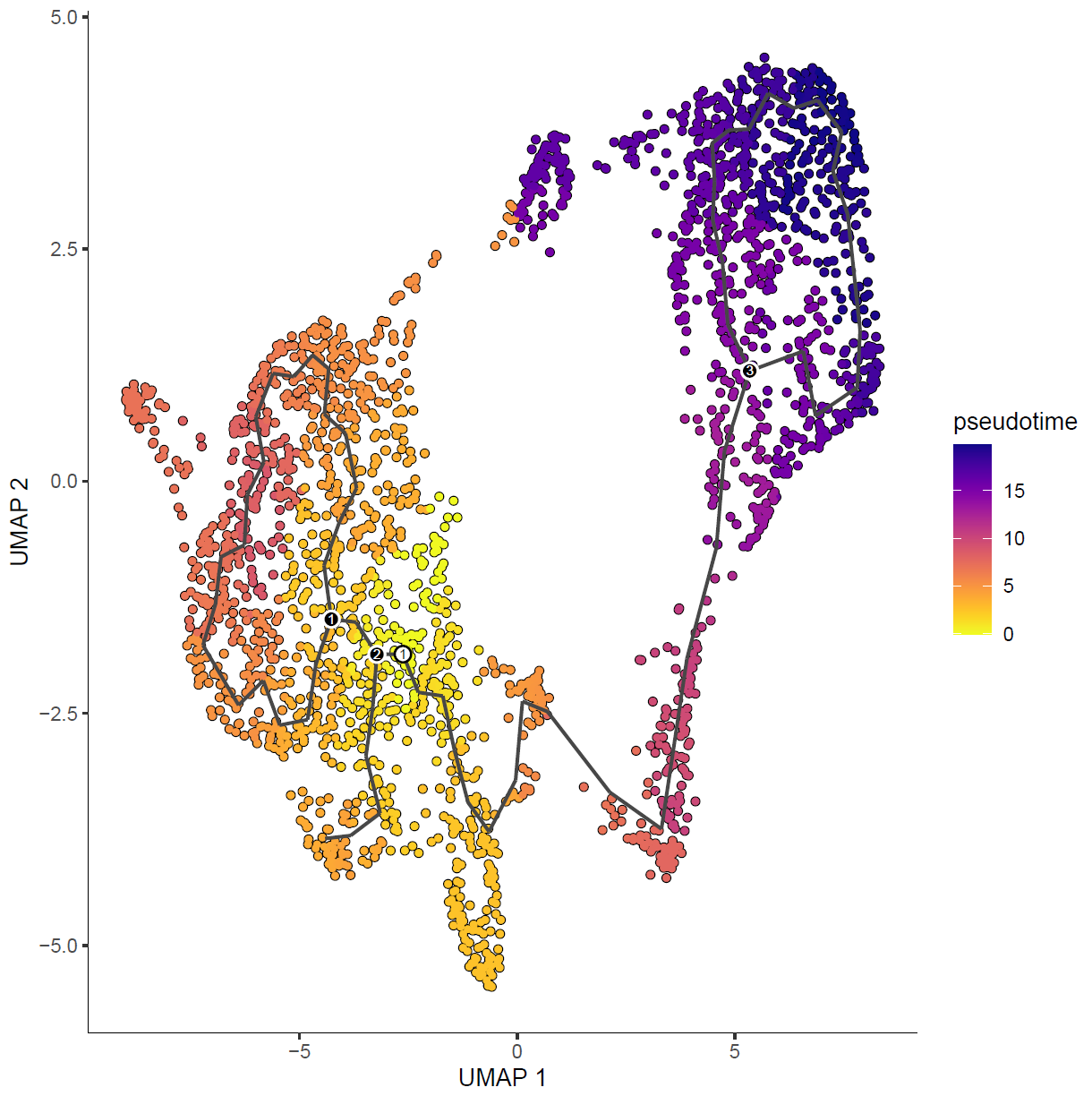

Figure S7 - Pseudotime trajectory (Monocle 2) for tumour cells in PDX 272. Trajectory begins in the adenocarcinoma component (left; yellow/orange) and continues into the neuroendocrine component (right; blue/purple).

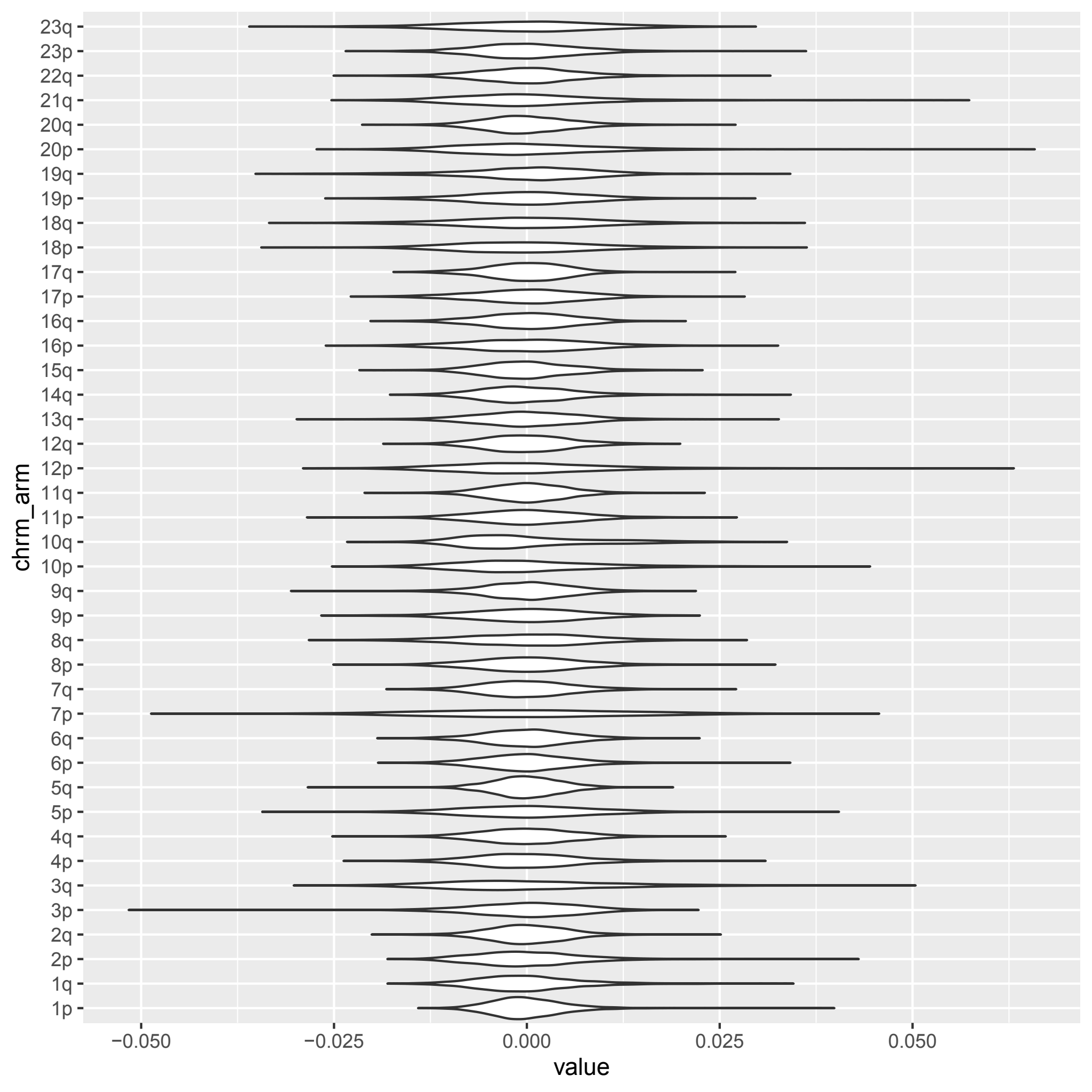

Figure S8 - Relative expression of genes on each chromosome arm for tumour cells from PDX 470B.

**Tables** (Provided as separate files)

Table S1 – Summary of single-cell RNA-sequencing data for each PDX tumour included in this study.

**Supplementary Note 1 – Benchmarking Data Integration Methods**

Three algorithms for integration of single-cell RNA-sequencing were evaluated: Seurat's canonical correlation analysis (CCA), LIGER's integrative non-negative matrix factorisation (iNMF) and Cluster Similarity Spectrum (CSS) from Simspec (Figure S9). CCA aims to identify matching cell pairs across datasets that are maximally correlated. These "anchors" represent a similar biological state, weighted based on the overlap in their nearest neighbours. This creates a reference to transfer data and metadata from one experiment to another. iNMF provides a low-dimensional space in which each cell is defined by one set of dataset-specific factors; each factor usually corresponds to a biologically interpretable signal representing a particular cell type. iNMF aims to identify shared and dataset-specific metagenes across datasets. CSS considers every cell cluster in each sample for integration as an intrinsic reference and represents each cell by its transcriptome similarities to clusters across samples. Details of how each algorithm was applied to the data are provided later in this Supplementary Note.

To benchmark and compare integration pipelines were benchmarked using data from two PDX models derived from MURAL patient 224 (Table S1). PDX 224R is a small cell NE mixed with adenocarcinoma grown in a testosterone supplemented with intact gonads, while PDX 224-Cx originated from grafting PDX 224R into a castrated host to generate a castration-resistant subline. We assessed the ability of each integration to correctly cluster together neuroendocrine cells from these two homologous PDX tumours from intact and castrate hosts while excluding the adenocarcinoma cells from PDX 224 into separate clusters. Four metrics of integration quality were used: silhouette coefficient, mixing metric, and local structure metric. Details of quality metric calculation are provided later in this Supplementary Note.

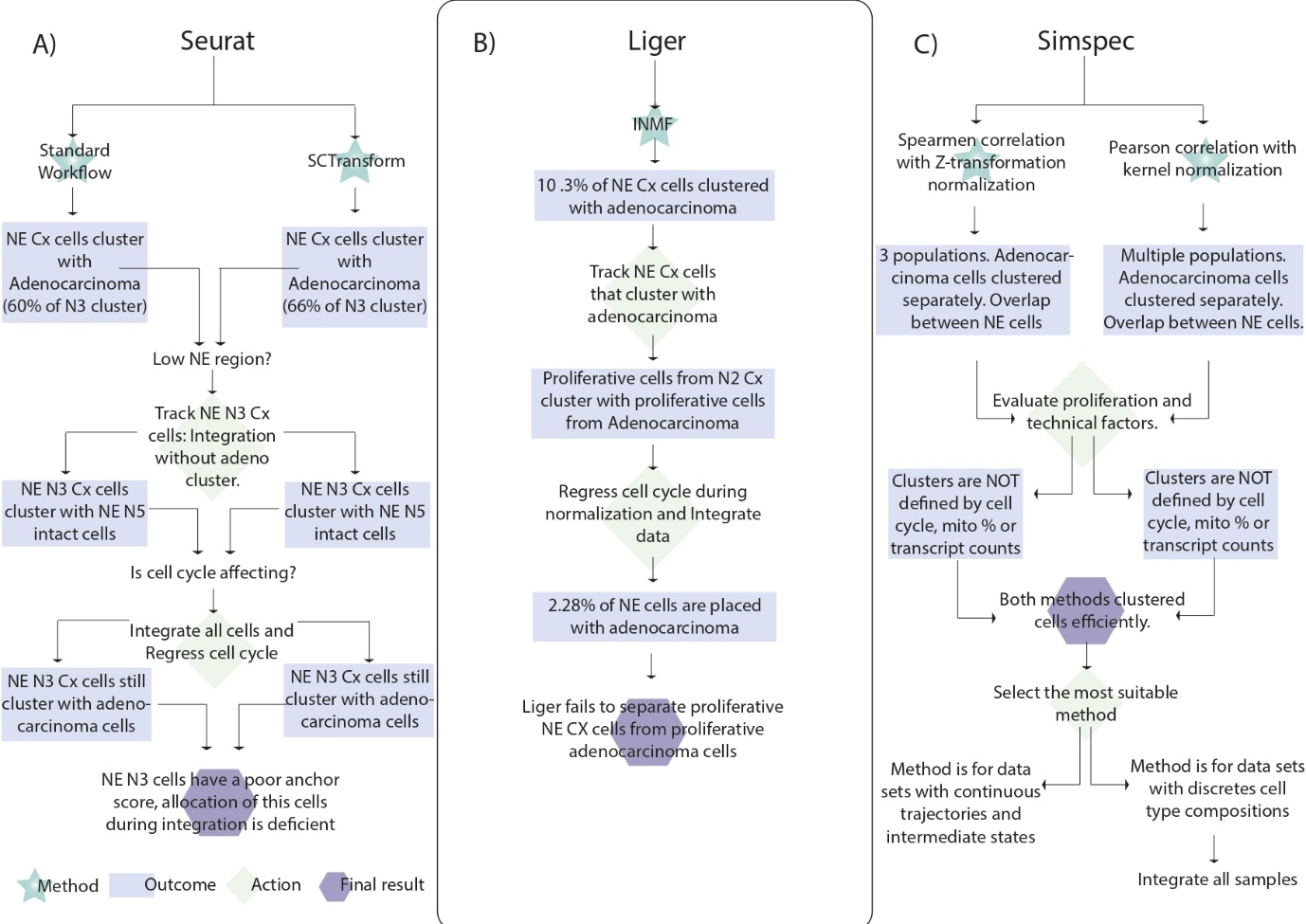

**Figure S9: Workflow chart of benchmarking process.** **A)** Seurat integration workflow process. Seurat has two ways of pre-processing the data for integration; standard workflow and SCTransfrom. Standard workflow includes normalization, finding variable features and scaling data. SCTransfrom is a wrapper that includes the standard workflow, but the data is not scaled. **B)** Liger integration workflow process. Integrative non-negative matrix factorization (iNMF) method was used. **C)** Simspec integration workflow process. The cluster similarity spectrum method was used. Two correlation methods might be used: Pearson or spearman. Pearson correlation uses kernel normalisation and Spearman correlation uses Z-transformation normalisation.

The CSS Simspec integration method was ultimately chosen due to its ability to segregate adenocarcinoma from neuroendocrine cells and overlapping common neuroendocrine populations while preserving transcriptional heterogeneity from each sample. Initial integration using Simspec showed the best separation between adenocarcinoma cells and neuroendocrine populations (Figure S10, panel C1). No neuroendocrine cells were detected in the adenocarcinoma clusters. Metrics showed that Simspec integration method was able to remove batch effect (AWS: 0.3, mix metric: 298) while preserving local structure of the clusters (score: 0.34). Entropy scores were low in all clusters, meaning that no adenocarcinoma cells were mixed with NE cells in any cluster and vice versa (Figure S11).

Further analysis was performed to ensure that the grouping was not based on any technical source of variation, such as transcript count and mitochondrial percentage. The mitochondrial percentage is less than 20%, and clusters are not formed by a high or low mitochondrial gene content (Figure S10, panel C2). Transcript count showed an even distribution of cells with ~3000 transcripts per cell (Figure S10, panel C3), proliferative cluster showed a higher transcript count of ~5,000 per cell, as expected (Figure S12B).

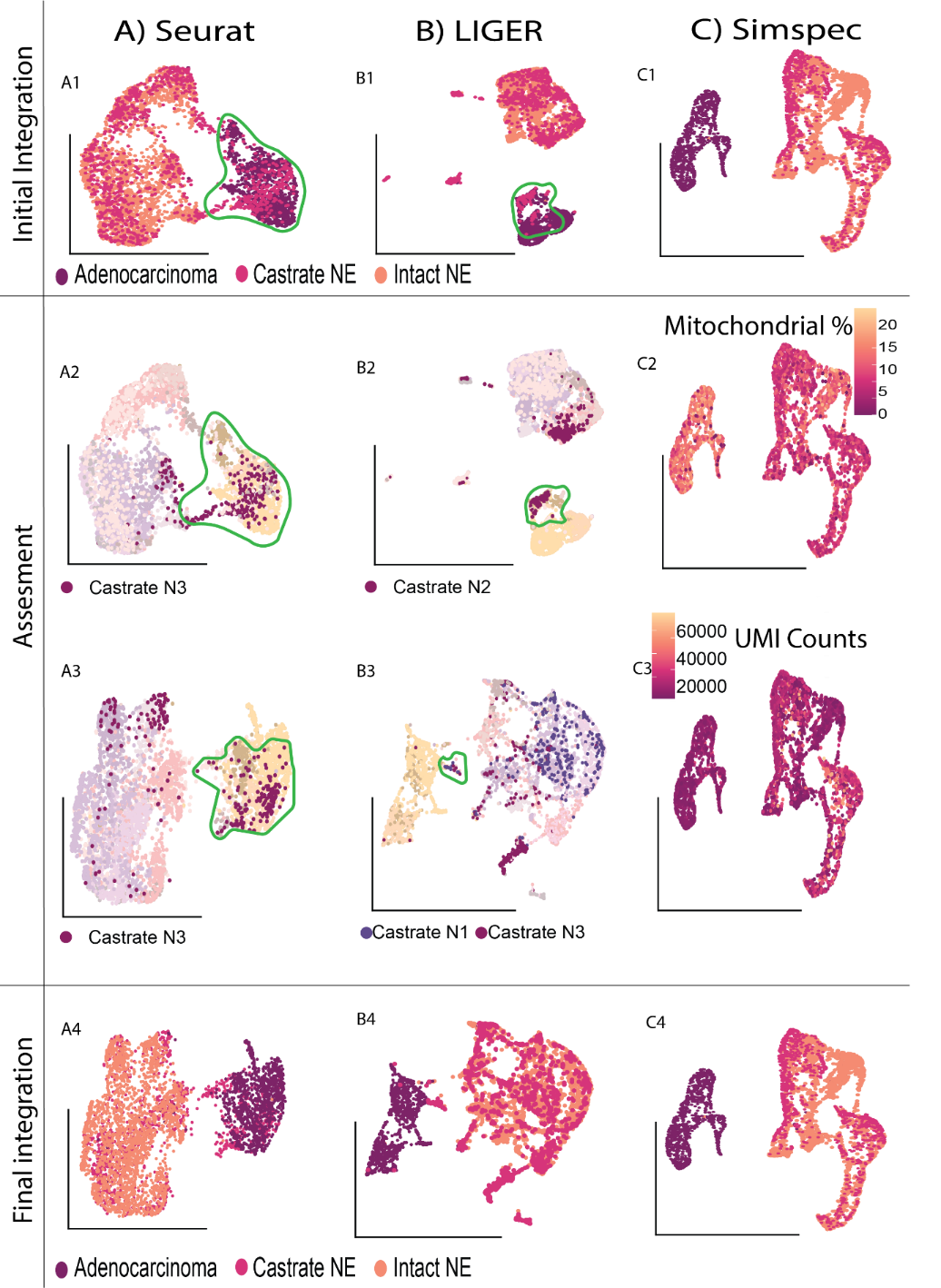

**Figure S10 Integration outcomes of 3 different pipelines**. UMAPS represents the outcome of each step of the process A) Seurat pipeline. A1) UMAP representing the Seurat integration using SCTransform as a normalisation method. In a green outline, castrated cells cluster with adenocarcinoma. A2) UMAP shows the main subpopulation of castrated cells that cluster with adenocarcinoma. A3) UMAP represents the integration after removing the cell cycle phase as a source of variation. Castrated cells from cluster N3 remain clustering with adenocarcinoma and are outlined in green. A4) UMAP represents the outcome of integration with Seurat. B) Liger pipeline. B1) UMAP representing the Liger integration. In a green outline, castrated cells cluster with adenocarcinoma. B2) UMAP shows the main subpopulation of castrated cells that cluster with adenocarcinoma—outlined in green castrated N2 cells. B3) UMAP represents the integration after removing the cell cycle as a source of variation. Castrated N1 and N3 cells clustered with adenocarcinoma, outlined in green. B4) UMAP showing the outcome of integration with Liger. C) Simspec CSS pipeline. C1) UMAP represents the initial outcome of the integration using Simspec. Circled in a green outline, a cluster is uniquely formed by castrated cells. C2) UMAP coloured by the expression of mitochondrial percentage. C3) UMAP coloured by the transcript (UMI) count. C4) UMAP with the Simspec integration pipeline after QC.

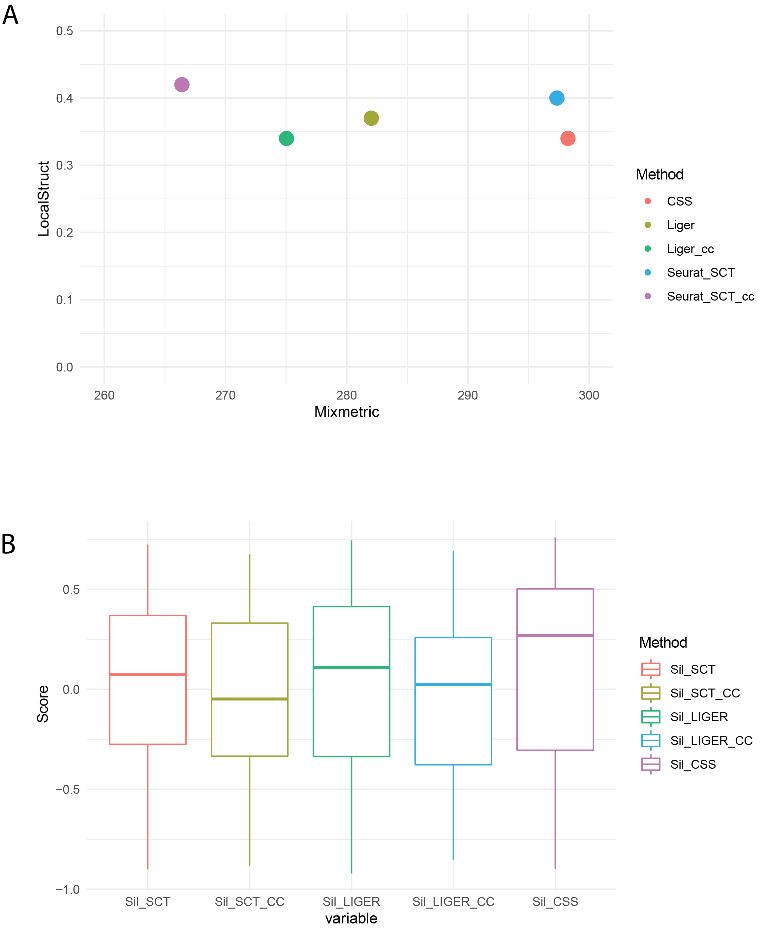

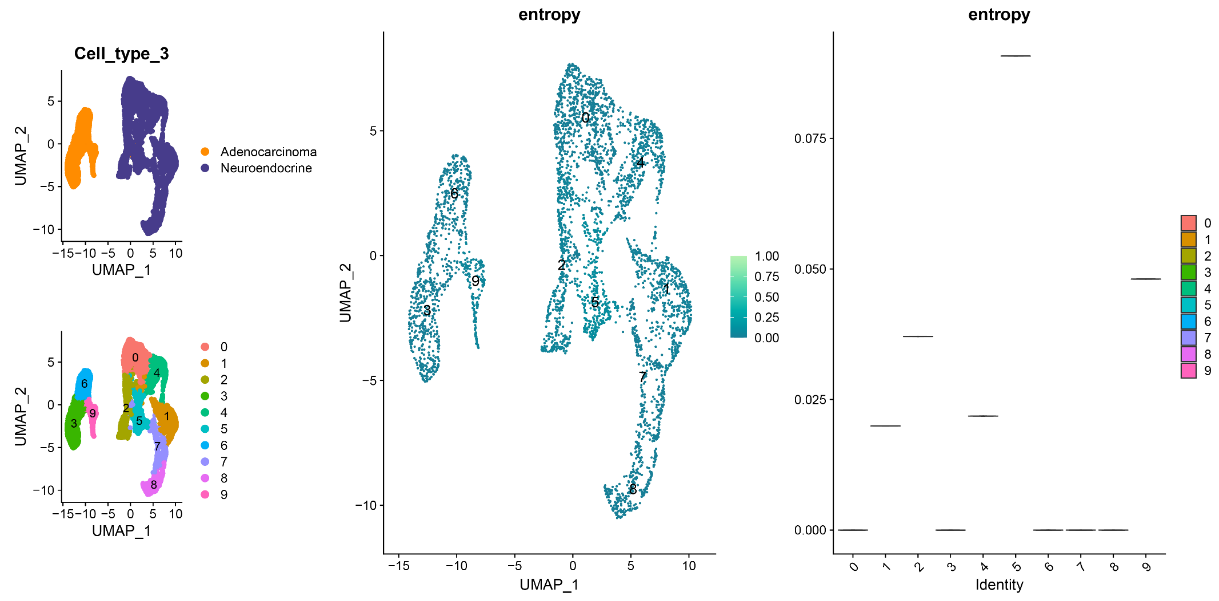

C

**Figure S11: Metrics for evaluating integration performance**. A) Local structure and Mix metric depicted in a dotplot. B) Boxplot of AWS scores per cell resulting from each integration method. C) UMAPs depicting cell type (Adenocarcinoma and neuroendocrine), clusters, and entropy score. Violin plot showing the score per cluster.

Furthermore, cancer signatures and differential gene expression analysis were evaluated on the integrated dataset to identify if heterogeneity was preserved (Figure S12 D-E). After integration, clusters with cancer signatures such as EMT, metastasis, angiogenesis, stemness and apoptosis were detected. Like cancer signatures, markers that previously characterised the heterogeneity of each tumour were preserved and detected across samples (Figure S12D). Clusters 0, 1 and 2 showed some pattern differences in the expression of specific markers, and this may be because of castration (Figure S12E).

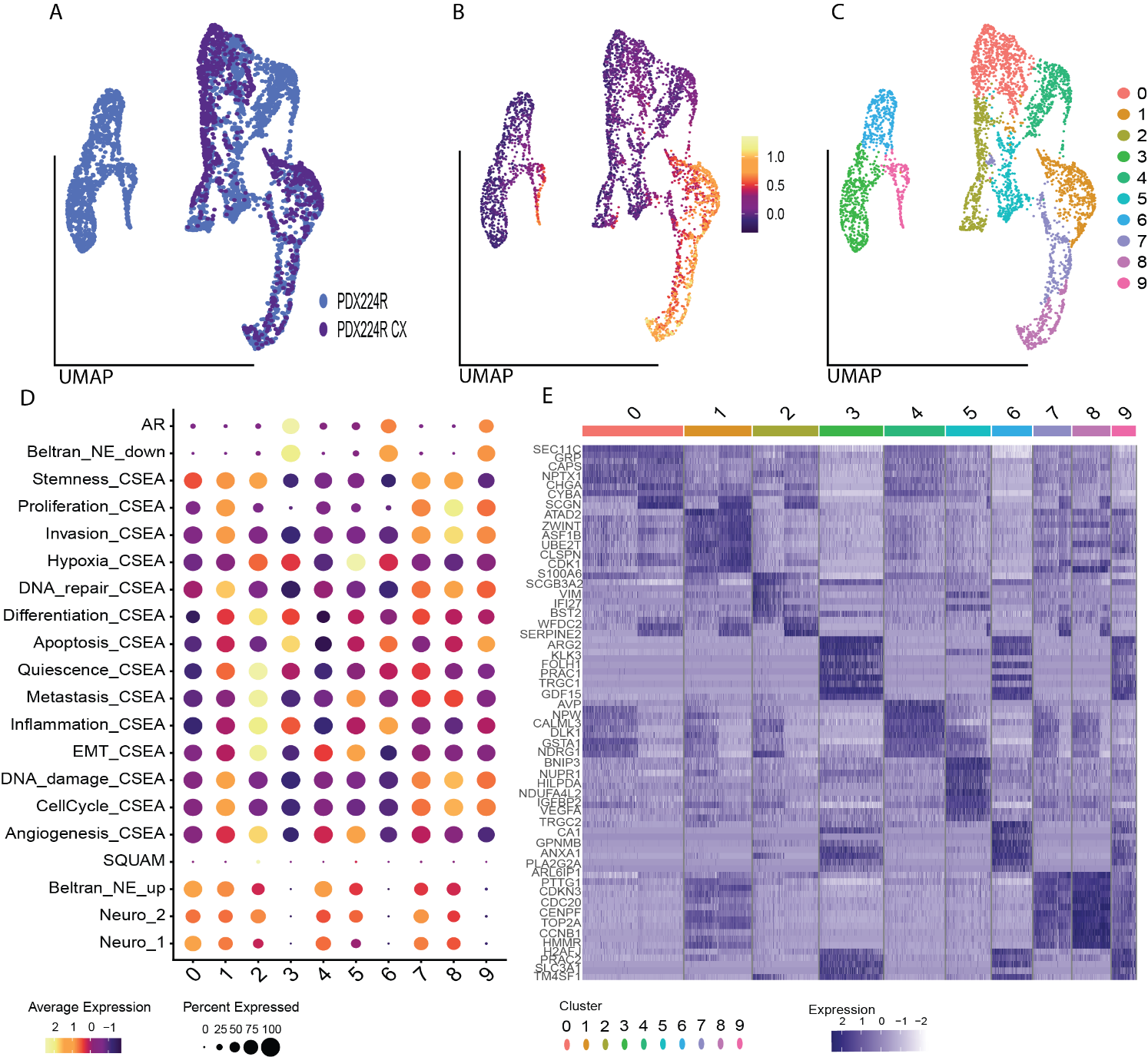

**Figure S12 Tumour heterogeneity is preserved after integration with CSS.** A) UMAP representing the integration of PDX224R and PDX 224R-Cx. B) UMAP depicting proliferative cells. A score close to one indicates a high expression of proliferative genes. C) UMAP showing the 10 clusters formed after integration. Clusters were defined with the same method used on previous chapter. D) Cancer CEA, NEURO I-II, Squamous and AR responsive signatures. The ratio of the circle represents the proportion of cells that express such signature, the colour of the circle represents the level of expression, yellow represents a high expression, purple represents a low expression. E) Heatmap showing the top markers per cluster

Implementation of integration algorithms

**Seurat SCTransform**

To integrate multiple datasets with Seurat, individual normalisation of each sample was done before integration. The normalisation function was “SCTransform”, based on regularised negative binomial regression. During normalisation, 3,000 variable features were selected for downstream analysis. After normalisation, a list of the objects to integrate was created. Next, setting features to use when integrating multiple samples was done using “SelectIntegrationFeatures”. This function generates a vector of the selected top scoring ranked features that will be used in the next step. Then, the function “PrepSCTIntegration” determines the features to use in the downstream integration procedure and ensures that sctransform residuals are present for all features that will be used as “anchors” in all datasets. The next step was to find features that could be used as anchors; the function “FindIntegrationAnchors” was used with standard parameters to create an anchor set object; this gene set will be used for integration. “IntegrateData” was then used to integrate all datasets given the previously pre-computed anchor set.

**Liger**

The “Seurat Wrappers” package was used to integrate Seurat objects with Liger. Before integrating, the count matrices of each sample must be merged into one Seurat Object; cell IDs were added, and the “merge” function from Seurat was used. The pre-processing steps include normalisation, finding variable features and scaling data. Normalisation was done with the “NormalizeData” function and finding variable features with “FindVariableFeatures”; both functions used default settings. To scale the data, the function “ScaleData” from Seurat was used; data was split by sample and not centred. The integration process starts with running the integrative non-negative matrix factorisation on the normalised and scaled datasets using “RunOptimizeALS”. The k parameter was set to 20, and the object was split by sample (processed separately); the rest of the parameters were used as default. The next step is to use the “RunQuantileNorm” function with the resulting factors to cluster cells jointly and perform quantile normalisation by dataset, factor, and cluster to integrate the datasets fully. Default parameters were used, and the object was split by sample. Louvain clustering was performed after integration using “FindNeighbours” and “FindClusters” from Seurat. The dimensional reduction was made using “RunUMAP” from the Seurat package.

**Simspec (CSS)**

A Seurat object is required to integrate single-cell data using the cluster similarity spectrum (CSS) algorithm in the Simspec package. The data was pre-processed with Seurat, and normalisation, finding variable features, scaling the data, PCA and dimensional reduction UMAP were needed before integration. The “cluster_sim_spectrum” function was used to integrate the data, the correlation method used was “Pearson”, and the spectrum type used was “corr_kernel”, cluster resolution was set at 0.3, and the label tag was defined as samples name. After integration dimensional reduction methods UMAP and PCA were run, the type of reduction used was “css” and “css_pca”, respectively, and ten dimensions were selected for each step. Then, the “FindNeighbors” and “FindClusters” functions were used to calculate clusters after integration. The resolution was set at 0.3, and 10 dimensions were used.

Metrics for evaluating integration performance

To compare the integration results, three measures of integration quality were used: the silhouette coefficient, a mixing metric, and a local structure metric.

*Silhouette coefficient*

The cluster R package (version 2.1.4) was used to calculate the silhouette coefficient. Before computing the silhouette score, distances were calculated using the code from Stuart, 2019 (1). The "Dist" function from the package Stats (version 3.6.2) in R was used for the distance matrix computation. Distances were computed using the UMAP space defined by the two dimensions for all methods. Labels of clusters must be defined; the previous ID of each cluster in each sample was used as a label. Then, the silhouette score was calculated using the distance matrices previously calculated and the integer vector with the previous cluster labels. Then the silhouette score was added to the metadata of the integrated Seurat object. A score of 1 or close to 1 represents dense and well-separated clusters. Furthermore, a score of 0 or -1 corresponds to overlapping clusters. A higher score equals high performance.

*Mixing metric*

Stuart et al. in 2019 designed the mixing metric (1). It is constructed to evaluate how well-mixed the input datasets were after integration. They reason that if neighbouring cells from a cell are well mixed, the closest neighbours should contain at least a small number (k=5) of cells from each dataset. If poorly mixed, the cell will be surrounded by a small subset of datasets or only the cells from their dataset. The function of "MixingMetric" in the Seurat R package was used. This function examines the local neighbourhood of each cell and determines for each group (clusters of datasets after integration) the k nearest neighbour and what rank that neighbour was in the overall neighbourhood. Then it subtracts the median across all groups as the mixing metric per cell. For this analysis, the grouping variable was the previous ID, the reduction used was "umap", the dimensions to use were 1:2, the k parameter was set to 5 and the max.k was set to 300; this was the suggested parameter by the user guide. For this metric, the higher the score, the better the integration.

*Local structure metric*

It is a metric designed to measure how well each dataset's original structure (cluster) is preserved after integration. The function "LocalStruct" splits the data back into its original dataset, re-calculates the PCA on the uncorrected data, and detects the k= 100 closest nearest neighbours. This function also computes the 100 nearest neighbours based on the PCA of the integrated dataset. Then, the intersection of these two neighbours for each cell is calculated, and the overlap fraction is computed. The higher the score, the best performance.

**Entropy**

To calculate the cluster sample diversity using Shannon Entropy, we used the Seurat R package (version 4.0.4) in R software. The function "calc_diversity" was employed, which takes a Seurat object as input. The Seurat object should have columns indicating sample IDs and cluster IDs. The argument "sample_id" specifies the name of the column containing the sample ID information, and "group_id" specifies the name of the column containing the cluster ID information. We defined the "sample_id" column as "orig.ident," which indicates the original identity of the samples, and "group_id" as "cell_type," which represents the cells that are labelled as adenocarcinoma or neuroendocrine.

The Shannon Entropy method was applied to calculate the diversity of clusters across samples. Shannon Entropy is a commonly used metric for measuring diversity in a dataset. It provides a quantitative measure of the uncertainty or randomness of a system, in this case, the distribution of clusters across samples.
